# The role of mental simulation in primate physical inference abilities

**DOI:** 10.1101/2021.01.14.426741

**Authors:** Rishi Rajalingham, Aida Piccato, Mehrdad Jazayeri

## Abstract

Primates can richly parse sensory inputs to infer latent information, and adjust their behavior accordingly. It has been hypothesized that such flexible inferences are aided by simulations of internal models of the external world. However, evidence supporting this hypothesis has been based on behavioral models that do not emulate neural computations. Here, we test this hypothesis by directly comparing the behavior of humans and monkeys in a ball interception task to that of recurrent neural network (RNN) models with or without the capacity to “simulate” the underlying latent variables. Humans and monkeys had strikingly similar behavioral patterns suggesting common underlying neural computations. Comparison between primates and a large class of RNNs revealed that only RNNs that were optimized to simulate the position of the ball were able to accurately capture key features of the behavior such as systematic biases in the inference process. These results are consistent with the hypothesis that primates use mental simulation to make flexible inferences. Moreover, our work highlights a general strategy for using model neural systems to test computational hypotheses of higher brain function.

## Introduction

From just a few glances, we can parse the structure of a novel scene, generate a rich understanding of its components, and use this understanding to make general inferences and predictions (Battaglia, Hamrick, and Tenenbaum 2013). This understanding helps us infer the latent states of objects and events, predict plausible and implausible future states, plan intervening actions, and anticipate the consequences of those actions. Despite the centrality of these capacities in human intelligence, the underlying computations remain unknown.

A dominant theory is that the brain constructs mental models of the physical world and relies on **mental simulations** of those models for making inferences (Craik 1952; Battaglia, Hamrick, and Tenenbaum 2013; Hegarty 2004). Mental simulations are also thought to underlie other cognitive functions such as imagination (Shepard and Metzler 1971; Hassabis, Kumaran, and Maguire 2007) and counterfactual reasoning (Gerstenberg and Tenenbaum 2017). Currently, the strongest evidence in support of the mental simulation hypothesis comes from comparing human behavior to predictions made by simulations of high-level computer programs that are analogous to how engineers run simulations of physical systems (Battaglia, Hamrick, and Tenenbaum 2013; Ullman et al. 2017). However, it is not known whether these observations reveal a fundamental aspect of the underlying brain algorithms or are a consequence of the class of cognitive models used to assess behavior. Indeed, many behaviors that are proposed to rely on mental simulations can also be accounted for by automatized functional approximations implemented by distributed neural network models (Lerer, Gross, and Fergus 2016). Therefore, the plausibility of the internal simulation hypothesis rests critically on whether or not the neural systems that support human inference also rely on simulation strategies (see Figure 1A).

**Figure 1.**
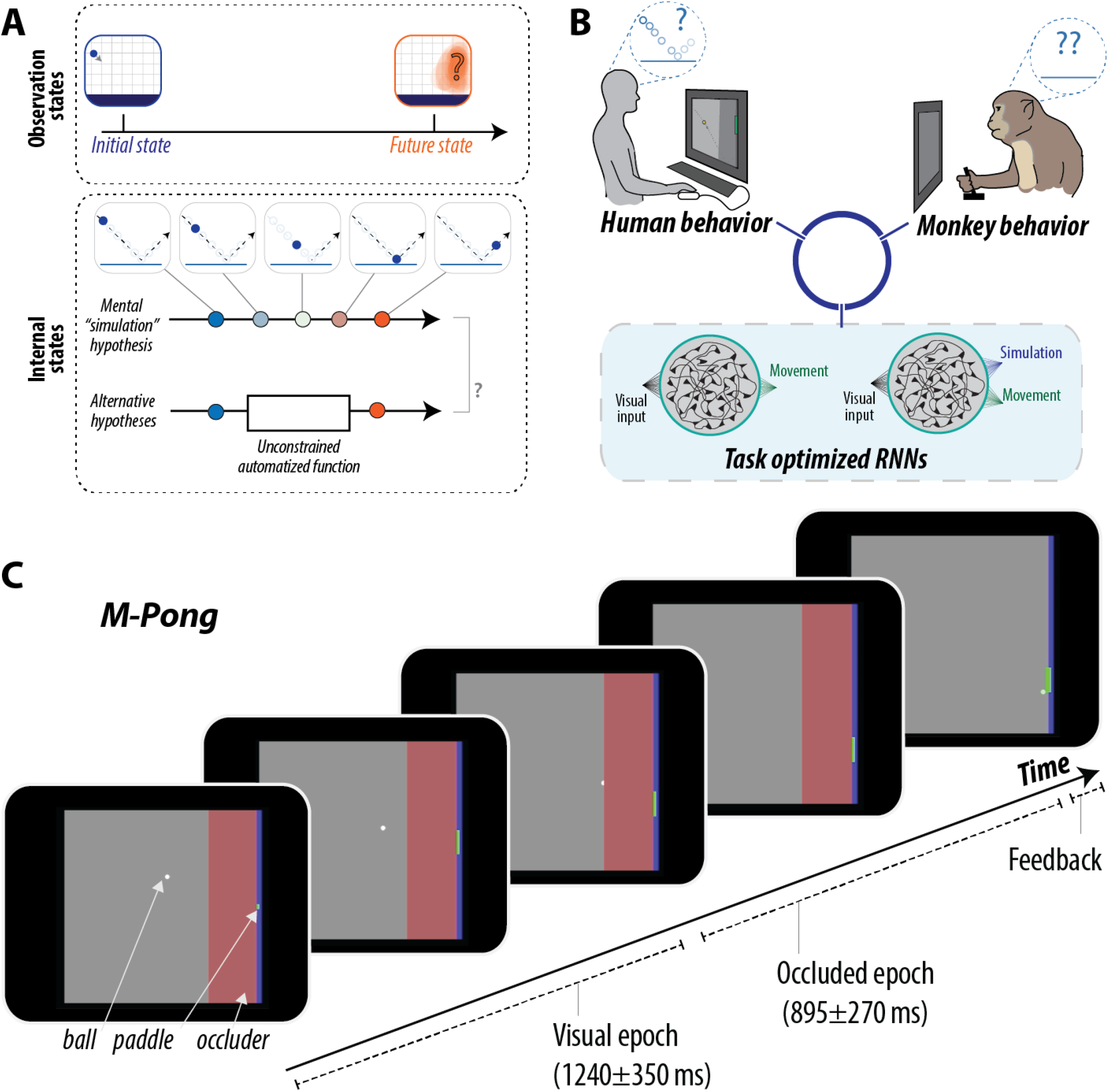
**(A) Conceptual schematic**. A dominant theory in cognitive science is that humans make inferences about external processes using internal simulations of the external physical world. To illustrate this, the top panel depicts observations of the external world, e.g. a ball falling towards the ground. In making an inference about the future state of the external world, do humans use a strategy based on “mental simulation” or not? In other words, do the internal states in the brain reflect the dynamics of the external world, or can they be explained by other complex, distributed functions? **(B) Experimental strategy.** We test this hypothesis by directly comparing the behavior of humans, monkeys and task-optimized recurrent neural network models. We trained RNN models to solve the same task as humans and monkeys (see panel C), but additionally optimized some models to perform proxies of “simulation.” **(C) Behavioral task.** We developed a behavioral task (*M-Pong*, reminiscent of the computer game called Pong) that aims to probe the ability of humans and monkeys to flexibly and rapidly predict the future state of a previously learned rich physical world. The objective of the task is to control the vertical position of a paddle to intercept a ball moving across a two-dimensional frame with reflecting walls. The frame additionally contained a large rectangular occluder right before the interception point such that the ball’s trajectory was visible only during the first portion of the trial. Subjects initiated each trial by fixating on a central fixation dot (not shown). Following this fixation acquisition, subjects were allowed to make any eye movements and freely view the screen, upon which the M-Pong frame was rendered onto the screen spanning 20 degrees of visual angle: the ball was rendered at its randomly sampled initial position, and the paddle was rendered in the central vertical position at the right edge of the frame. The paddle was initially rendered as a small, transparent green square, but turned into a full paddle when the subject first initiated paddle movements (i.e. pressed a key) to enforce that subjects performed a movement on all trials (see Methods). On every trial, subjects could move the paddle as soon as the ball began to move, and could drive it freely and at a constant speed in up or down directions from its initial position at the middle of the screen with the goal of intercepting the ball when it reached the paddle. Ball movement parameters (position, heading, speed) were randomly sampled on every trial. Critically, the visual input is completely uninformative once the ball goes behind the occluder; the M-Pong task forces subjects to make inferences about the dynamics of an unobserved external process.

To tackle this question, we took a rigorous computational approach involving detailed analysis of behavior in humans, monkeys, and a large battery of recurrent neural network (RNN) models with or without “mental simulation” capacities. We developed a behavioral task that demanded rapid learning and flexible generalization, and verified that both humans and monkeys (hereafter, “primates”) were able to perform the task. Having established a platform for testing the mental simulation hypothesis, we reasoned that if primates rely on simulation-like computations, then RNNs bestowed with the capacity to “simulate” should capture primates’ behavior more accurately than models that are not. We defined simulation in RNNs operationally as the ability to track latent environmental states in real time and in the absence of sensory inputs. Accordingly, we created a large collection of task-optimized RNNs with different types of inputs and internal architectures and asked whether those that were additionally optimized to perform simulations were better at explaining characteristic behavioral patterns in primates (Figure 1B). We found that the behavioral similarity of the RNNs to primates was specifically and systematically related to the degree to which RNNs faithfully simulate environmental states. Further analysis revealed how structured dynamic representations in the primate-like RNNs serve as a computational substrate for performing simulations.

## Results

### Behavioral task

We devised a behavioral task in which subjects had to control the vertical position of a paddle to intercept a ball moving across a two-dimensional frame with reflecting walls (Figure 1C). The ball’s initial position and velocity (speed and heading) were randomly sampled on every trial (Figure S1). Moreover, the frame contained a large rectangular occluder before the interception point such that the ball’s trajectory was visible only during the first portion of the trial. Depending on the ball’s initial position and velocity, the visible and occluded portions of each trial lasted 1240+/−350 and 895+/−200 ms, respectively. Subjects initiated each trial by fixating on a central fixation dot, but were subsequently free to make any eye movements. On every trial, subjects could move the paddle as soon as the ball began to move, and could drive it freely and at a constant speed in up or down directions from its initial position at the middle of the screen with the goal of intercepting the ball when it reached the paddle (Figure 2A, left). We used identical task parameters and contingencies for humans and monkeys with two exceptions. First, humans moved the paddle using a keyboard, whereas monkeys used a one-degree-of-freedom joystick. Second, while both monkeys and humans received visual feedback at the end of each trial, monkeys additionally received a small juice reward when they successfully intercepted the ball. We refer to this task as *M-Pong* because of its similarity to the computer game Pong, and because of the presence of the occluder that necessitates mental (as opposed to visual) computations.

**Figure 2.**
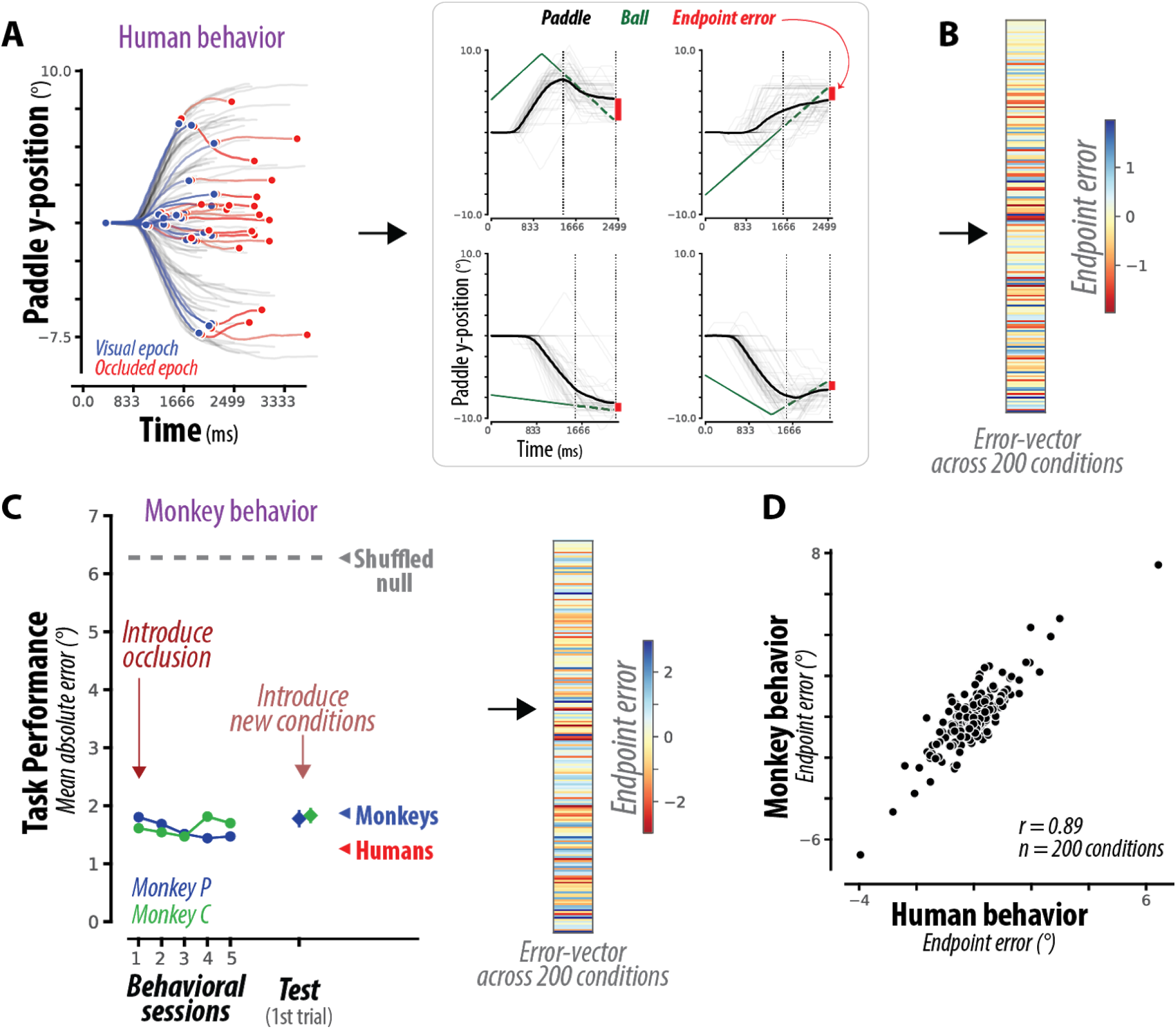
**(A) Human behavior**. (left) Each curve corresponds to the paddle position (in units of degrees of visual angle) over time for a single task condition, averaged across dozens of trial repetitions, separated into movements while the ball was visible (blue) or occluded (red). (right) For four example task conditions, this average paddle trajectory is overlaid on each individual trial repetition (light gray), as well as the true vertical position of the ball (green), from which the average endpoint error (red) is estimated. **(B)** Across 200 different task conditions, we measure a pattern of endpoint errors (termed error-vector) to characterize behavior. The error-vector is shown as a colored vector, with colors spanning mean ± 2SDs of the error range. **(C) Monkey behavior.** (left) After training monkeys to manipulate the joystick in order to control a paddle and intercept moving balls, we introduced the occluder, interleaving trials where the ball was completely versus partially occluded. Monkeys reached high performance under occlusion on the very first behavioral session, and maintained this high performance over subsequent sessions (quantified with the mean absolute error, in units of degrees of visual angle). Furthermore, monkeys maintained high performance when tested on 150 novel task conditions, and this generalization was immediate, with comparable high performance on the very first trial of a new condition for both monkeys (“test” marker, mean + SE across 150 conditions). For comparison, chance performance (shuffled null) is shown via the gray dashed line and arrowhead, and the final performance of humans and monkeys is shown via red and blue arrowheads respectively. (right) Following training, we characterized monkey behavior across 200 task conditions with a pattern of endpoint errors, shown as a colored vector, with colors spanning mean ± 2SDs of the error range. **(D) Comparison of human and monkey behavior.** The scatter shows the average endpoint error for each of the 200 conditions, for humans versus monkeys. We observe a remarkable similarity in the error-vectors of humans and monkeys.

M-Pong has several desirable features. First, unlike tasks that rely on repeated measurements for a small set of conditions, in M-Pong, nearly every trial is unique (see Methods). This property makes it virtually impossible to adopt a model-free stimulus-response policy; instead, subjects have to form an internal model of the task that can be used to generalize across new parameters and conditions. Second, unlike simple working memory tasks, the appropriate response in M-Pong is a time-varying and non-linear function of previously observed stimuli. This property places a powerful constraint on the space of models that can capture the behavior. Third, the presence of the occluder motivates an online estimation strategy to predict the position of the ball at the time of interception despite the absence of any visual input. In this regard, a plausible strategy is to mentally simulate the physical movement of the ball as it moves behind the occluder. Note however that adopting a simulation strategy is not obligatory; since the initial position and velocity of the ball fully determine all its future states, a sufficiently complex nonlinear function would be able to solve the task without any simulation. Fourth, as we will show, monkeys were able to learn this task remarkably fast and were able to effortlessly generalize.

### Comparing human and monkey behavior

We collected behavioral data from 12 humans performing 200 unique task conditions (i.e., different initial ball position and velocity), all randomly interleaved. For all subjects, we measured the position of the paddle throughout the trial (Figure 2A, left). Subjects began moving the paddle early in the trial while the ball was still visible (Figure S2F), and generated smooth movement trajectories that continued throughout the occluded portion of the trial (Figure S2F).

We quantified overall performance using the mean absolute error (MAE), computed as the absolute difference between the final ball position and the final paddle position, averaged across all trials and all conditions. Importantly, we went beyond the overall performance metric and additionally analyzed condition-specific errors. For each of the 200 unique conditions, we quantified the average endpoint error; i.e. the signed difference between the final ball position and the final paddle position, averaged across trial repetitions of a specific condition (Figure 2A, right). This resulted in a 200-dimensional *error-vector* that we used to compare behavior across humans, monkeys and RNNs.

The analysis of the error-vector in humans led to two important conclusions. First, the error-vector was highly similar across humans (see Figure S2B) indicating that humans solve the task using a similar inference strategy. Second, the pattern of errors could not be straightforwardly explained in terms of the ball’s initial position and velocity (Figure S2D). From these observations, we concluded that the common pattern of errors across humans reflects the inference strategy they rely on to solve the task, and can thus serve as a metric to compare with monkeys and RNNs.

Next, we trained two monkeys to perform the M-Pong task. After an initial familiarization with the joystick as the means for controlling the paddle, monkeys were trained to intercept a moving ball, without any occluders, which they mastered after several days. Next, we introduced the occluder, interleaving trials where the ball was either completely or partially invisible. To our surprise, monkeys reached high performance under occlusion on the very first behavioral session, and maintained this high performance over subsequent sessions. This is shown in Figure 2C, which shows that the per-session error of monkeys (green/blue circles) is dramatically lower than chance performance (shuffled null, gray dashed line and arrowhead), and approximately matches the final performance of humans and monkeys (red/blue arrowheads). This observation suggests that the computational demands of M-Pong are broadly compatible with macaque monkeys’ inductive biases.

Monkeys were trained on the same exact unique 200 conditions as humans. However, we used a curriculum that allowed us to test the monkeys’ ability to generalize. We first introduced 50 of the unique conditions, and later used the remaining 150 conditions as a test for generalization. Both monkeys were able to effortlessly generalize, and maintain the same performance level on the very first trial of the test conditions (Figure 2C, “test” on abscissa). These results indicate that monkeys solve M-Pong using a internal model that can be deployed flexibly without over-training or adopting stimulus-response memorization strategies.

Next, we analyzed monkeys’ behavior based on the systematic pattern of errors across the 200 unique conditions. We first verified that error patterns were similar between the two monkeys (Figure S3B), and could not be explained in terms of the ball’s initial position and velocity (Figure S3D). Next, we compared the error-vector between monkeys and humans. The conditions that monkeys found to be particularly difficult were similarly difficult for humans (Figure S2C, S3C). Moreover, the overall patterns of errors across the entire set of 200 unique conditions were highly similar between humans and monkeys (Figure 2D). Based on these observations, we concluded that monkeys rely on an inference strategy similar to humans, and that the pattern of errors across primates reflect this common strategy.

To quantify the strength of the similarity between a subject’s behavior and human behavior, we developed a summary statistic, which we term human-consistency score. We defined human-consistency score as the degree to which an error-vector was correlated with the error-vector derived from the behavioral responses in humans. To improve our estimate of human-consistency, the correlation coefficients were adjusted for sampling noise (see Methods). Moreover, to avoid overestimating human-consistency, errors were computed as residuals rather than simple differences, equivalent to computing a partial correlation between the paddle endpoint across conditions, accounting for the co-varying pattern of ground truth positions (see Methods).

Defined in this way, a human-consistency score of 1 would correspond to identical error patterns and a score of zero, to random errors. Human consistency score for monkeys was large (0.89 ± 0.003) but smaller than the ceiling value estimated based on a comparison of behavior between humans (0.95 ± 0.012, see Methods).

### Comparing primates and recurrent neural network models

RNNs have vast computational capacities and can, in principle, be trained to establish arbitrarily complex functions (Funahashi and Nakamura 1993). The RNN solutions one can find for a specific task are not unique, and can vary with various factors, including network architecture, training protocol, and perhaps most importantly, the optimization constraints one imposes on the solution(Maheswaranathan, Williams, and Golub 2019). We exploited this non-uniqueness property to ask whether RNNs of various architectures that are optimized to perform M-Pong would behave more similarly to primates if they were additionally optimized to simulate the ball position (Figure 3A).

**Figure 3.**
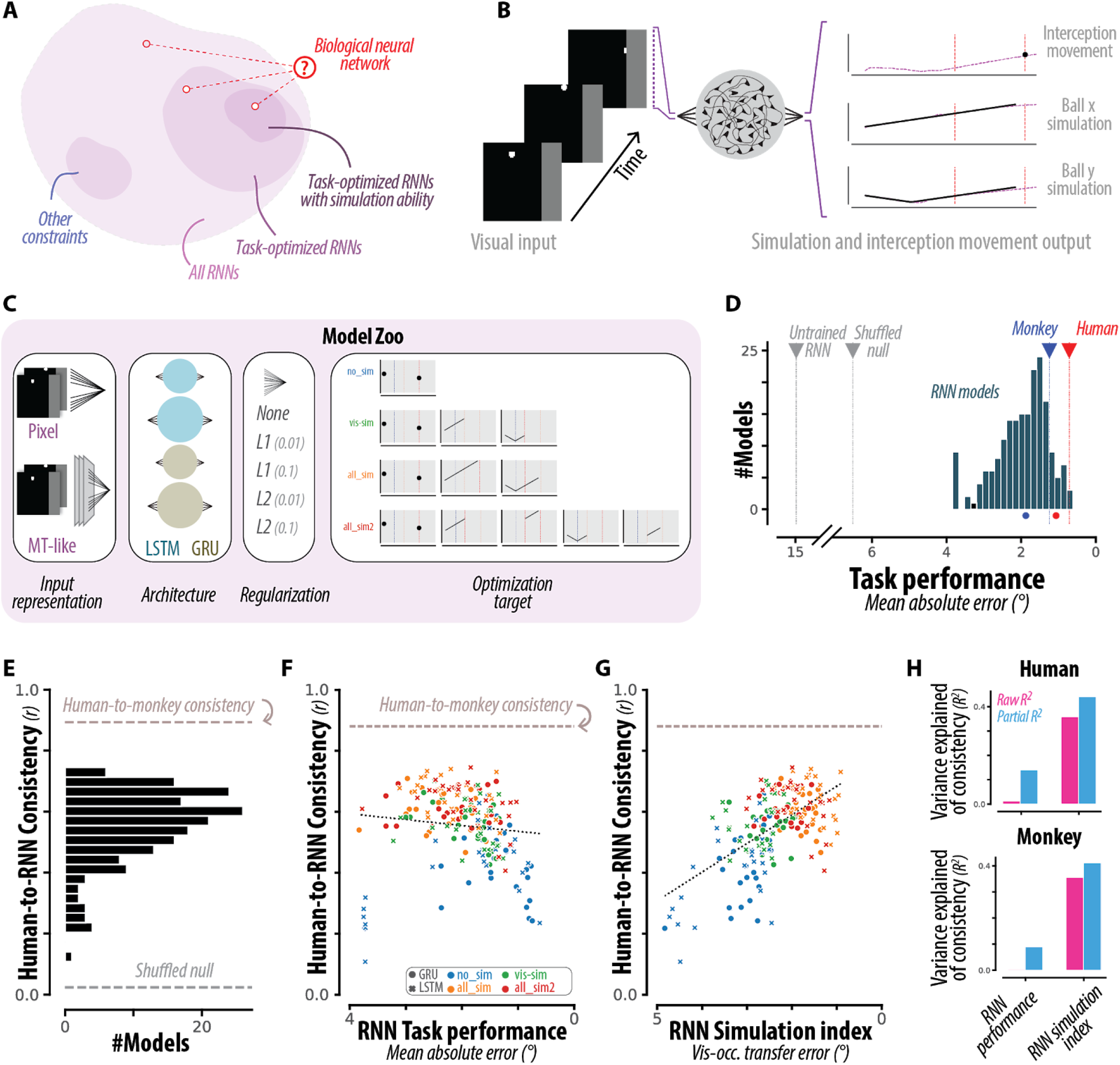
**(A) Conceptual schematic**. We reasoned that if flexible primate behavior in this task relies on simulation-like computations, then including specific functional constraints (e.g. the ability to “simulate”) on RNNs should lead to finding more brain-like algorithms. **(B) RNN behavior.** We trained several hundred RNNs to map a series of visual inputs (pixel frames) to a movement output, where the target movement output (black dot) corresponded to a prediction of the particular paddle position at a particular time point in order to intercept the ball. Some RNNs were additionally optimized to predict the position of the ball throughout the trial (black curves). **(C) Full set of RNN hyperparameter choices.** Different RNN models varied with respect to architectural parameters (e.g. different cell types, number of cells, regularization types, input representation types, etc.), and were differently optimized (either with or without internal simulation). Critically, RNNs were not optimized to reproduce primate behavior, only to solve the task. **(D) RNN performance.** RNNs were able to achieve high performance on this task, with accuracies comparable to humans and monkeys. Human/monkey performance is shown both with (colored circles) and without (colored triangles) the inclusion of errors stemming from trial-by-trial variability. Note that the abscissa is flipped such that left-to-right corresponds with increasing performance (i.e., decreasing error). **(E) Comparing human and RNN behavior.** We observed a broad distribution of human-consistency values, with some RNNs approaching human-monkey consistency (dashed line). **(F) Functional correlates of human-consistency.** Across all RNNs, scatter of human-consistency versus task performance (left) and simulation index (right). The variation in human-consistency across different RNNs did not strongly depend on overall task performance, with the behavioral patterns of the highest-performing RNNs typically diverging from those of humans. Instead, human-consistency was strongly correlated with a “simulation index”, an error metric of the ability to predict unobservable ball position. Note that the abscissas are flipped such that left-to-right corresponds with increasing performance (i.e., decreasing error) and increasing simulation ability (i.e., decreasing simulation error). **(G) Quantification of functional correlates.** The strength of dependence between the tested functional attributes and consistency with human behavior (top) and monkey behavior (bottom) is shown as a proportion of variance explained (R2, pink bars). Partial R2 (blue bars) measures this strength after accounting for covariations due to the other attribute.

We built each RNN as an input-output system that receives trial-by-trial sensory information about the ball and uses a linear readout to generate a scalar output to drive the paddle (Figure 3B). The database of RNNs we considered varied along several dimensions (Figure 3C) including cell type (LSTMs versus GRUs), number of cells (10 vs 20), input representation (pixel information only versus pixel information plus direction of motion), and regularization strategy (L1 versus L2 loss term). Consistent with the goal of the M-Pong task, we trained all networks using a standard performance-optimizing cost function to find solutions that minimize the error between the paddle and ball position along the y-axis at the time of interception with no constraint on how the paddle ought to move throughout the trial.

To examine the behavior of networks with simulation capacities, we augmented the cost function in subsets of networks by requiring the output of additional linear readouts to carry an explicit online estimate of the ball’s x and y position (Figure 3B,C). For each network architecture, we included four optimization strategies: 1) RNNs that were only optimized for the final paddle position without online simulation (“no_sim”), 2) RNNs that were additionally optimized to simulate ball position when the ball was visible (“vis_sim”), 3) RNNs that were optimized to simulate ball position throughout the trial (“all_sim”), and 4) RNNs that used different readout channels to simulate the ball position for the visible and occluded portions (“all_sim2”). Critically, none of our optimization factors were designed to make RNNs reproduce primate behavior. Instead, our goal was to test if any of the additional cost functions associated with simulation would enable RNNs of various architectures to spontaneously behave more similarly to primates.

We first verified that RNNs with different architectures and optimization choices were able to learn the task, and some were able to achieve performance levels comparable to humans and monkeys (Figure 3D). Next, we analyzed the entire zoo of RNNs in terms of the similarity of their behavior to that of the primates. To quantify the degree of similarity, we used the condition-dependent average-endpoint-error that we previously used to compare humans to monkeys. Human-consistency scores across RNNs were distributed broadly and were generally below the values we found for monkeys (Figure 3E).

We exploited the breadth of this distribution to ask what factors make certain RNNs behave more or less like primates. As a first step, we asked whether human-consistency of RNNs could be explained by their overall performance, which is a common observation in network models of vision and audition (Yamins et al. 2014; Kell et al. 2018). Results revealed no significant relationship between human-consistency and performance (*R^2^ = 0.01* and *0.00002, p > 0.05* for consistency to humans and monkeys, respectively, Figure 3F). This result indicates that the overall performance cannot reveal the factors that make RNNs more or less similar to humans.

Next, we focused on our primary objective of using our large RNN model zoo to test the role of simulation in M-Pong. Specifically, we asked whether networks that were optimized for simulating the ball position had higher human-consistency scores. To compare all RNNs on the same footing, we developed a “simulation index” that could be used in all networks, and would quantify the degree to which a network carries explicit information about the instantaneous position of the ball behind the occluder. The simulation index was computed as the mean absolute error between the true time-varying ball position and the predicted position from a cross-validated linear decoder (see Methods). Results revealed a strong relationship between simulation index and human consistency scores across the RNNs (Figure 3G). The simulation index was able to explain a large portion of the variance across humans and monkeys independently (*R^2^= 0.36* and *0.39, p < 1e-21* and *1e-23*, for consistency to humans and monkeys, respectively), and the effect was even stronger when we accounted for covariations due to performance (*R^2^=0.44, 0.43*, for consistency to humans, monkeys respectively Figure 3H), and greater than the corresponding estimates for task performance (*p < 1e-3, 1e-4*; dependent t-test of correlation coefficients). In sum, RNNs that carried explicit information about the latent position of the ball behind the occluder (i.e., performed “mental simulation”) were able to capture primate behavioral patterns more accurately than those that did not.

### Dynamics underlying primate-like behavior

We have shown that RNN models that simulate the ball position can accurately capture the behavioral error patterns of primates. We next asked *how* this optimization for simulation might lead to networks with more primate-like behavior. Given that optimization influences the network behavior by acting via the intermediary of its internal representations (see Figure 4A), we characterized the structure of RNN internal representations and related these characteristics to their consistency to primate behavior. We focused on dimensionality, slowness, and geometric curvature – three attributes that can be quantified in biological networks and can thus serve as predictions for future experiments on the primate brain.

**Figure 4.**
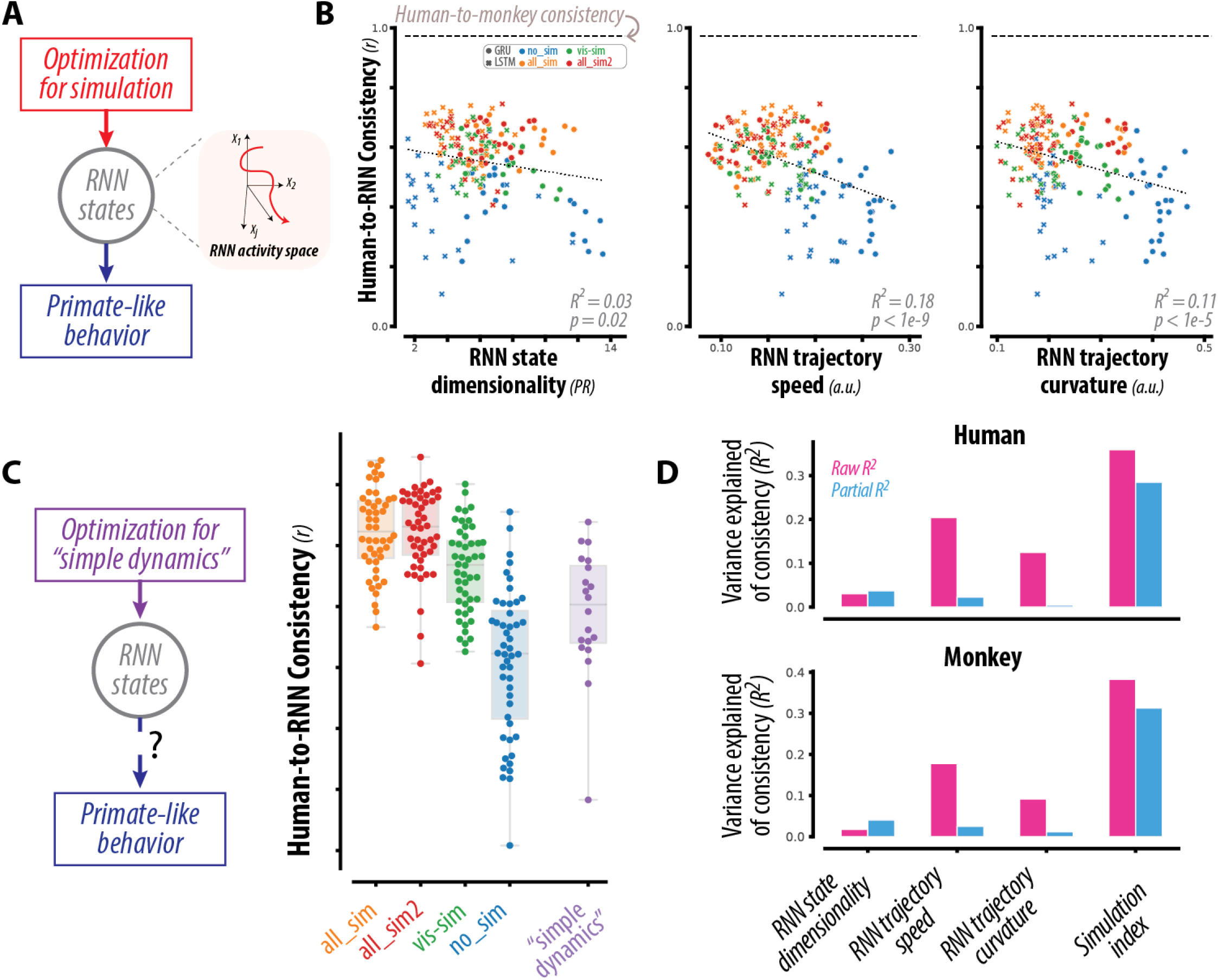
**(A)** Optimization for simulation ability influences the network behavior by acting via the intermediary of its internal representations. We characterized the structure of the internal representations of RNNs via specific geometric metrics. **(B)** Each panel shows the scatter over all RNN models of a representational attribute (dimensionality, speed, and curvature) against human-consistency. RNNs with simple dynamics (i.e. slow, smooth and low dimensional representations) exhibited more human-like behavior (and monkey-like behavior, not shown). **(C)** (left) To assess whether simple dynamics are sufficient to capture primate-like behavior, we constructed new RNN models that were identical to the simulation-optimized models, except for optimization consisting of task performance and specific regularization terms to favor slow and smooth dynamics. (right) Distribution of human-consistency scores for all RNNs, grouped by optimization types; the swarm plot shows individual models, and the boxplot shows the median, 1st and 3rd quartiles, and range of each disrtibution. RNNs optimized for simple dynamics better matched human behavior than RNNs optimized for task performance alone (blue vs purple distributions), but failed to capture primate behavior as well as models optimized for simulation (orange vs purple distributions). **(D)** The strength of dependence between the tested representational attributes and consistency with human behavior (top) and monkey behavior (bottom) is shown as a proportion of variance explained (R2, pink bars). Partial R2 (blue bars) measures this strength after accounting for covariations due to the other attribute. We found that simulation ability (quantified via the simulation index) better predicted consistency scores with respect to both human and monkey behavior.

Interestingly, we found that all three attributes were related to human-consistency scores across RNNs. Networks that exhibited “simple dynamics” – i.e. whose activity representations were lower dimensional, lower speed, and lower curvature – better predicted behavioral patterns of humans (Figure 4B). We quantified the relationship between these attributes and consistency scores by measuring the proportion of variance (*R^2^*) each feature could explain about consistency scores. Together, the three attributes predicted a modest proportion of variance associated with consistency scores (*R^2^ = 0.18*, and *0.16*, *p < 1e-10* and *1e-9*; with respect to human and monkey respectively). In other words, the state dynamics were generally smoother for RNNs that exhibited more primate-like behavior.

This observation raises the possibility that “simple dynamics” is sufficient for RNNs to emulate primate-like behavior. To address the possibility, we constructed new RNN models (termed “simple_dynamics”) that were identical to the simulation-optimized models in all respects except for their optimization, which consisted of both task performance and specific regularization terms to favor slow and smooth dynamics (see Figure 4C left, Figure S5A, Methods). As shown in Figure 4C (right), the consistency scores of these models were significantly improved relative to models optimized for task performance alone (*p < 1e-2, 1e-2*; t-test comparing “mov” vs “simple_dynamics” models for consistency to human and monkey behavior, respectively). However, they failed to capture primate behavior as well as models optimized for simulation (Figure 4C; *p < 1e-7, 1e-8*; t-test comparing “all_sim2” vs “simple_dynamics” models for consistency to human and monkey behavior, respectively).

To quantify the relative importance of “simple dynamics” versus “simulation ability” on consistency scores, we measured the proportion of variance (*R^2^*) of consistency scores that could be explained by each attribute, over all RNN models. We found that simulation ability (quantified via the simulation index) best predicted consistency scores with respect to both human and monkey behavior (Figure 4D, pink bars; *R^2^=0.35, 0.38, p < 1e-21, 1e-23,* for consistency with respect to human and monkey behavior respectively). This effect could not be explained by covariations due to simple dynamics, as evidenced by significant partial *R^2^* – the proportion of variance in consistency scores that could be uniquely explained by simulation ability (Figure 4D, blue bars; *R^2^=0.28, 0.31* for consistency with respect to human and monkey behavior respectively; *p < 0.005* for comparisons to all three dynamics attributes, dependent t-test of correlation coefficients).

Taken together, these results suggest that optimization for simulation drives RNNs to learn specific activity representations characterized by both simple dynamics (low dimensionality, speed, and curvature) as well as explicit task representations (i.e. linear projections of RNN states matching the ball position). These representations are characteristic of RNN models that exhibit primate-like behavior, and may be analogous to ones found in the primate brain.

### Dynamics underlying computations performed by RNNs

So far, we have treated RNNs as input-output “black box” instantiations of specific cognitive hypotheses. However, several recent studies (Mante et al. 2013; Chaisangmongkon et al. 2017; Sussillo et al. 2015; Russo et al. 2018; Wang et al. 2018; Remington et al. 2018; Sohn et al. 2019; Michaels, Dann, and Scherberger 2016) have shown the utility of reverse-engineering RNNs (Sussillo and Barak 2013) to shed light on how biological neural networks might perform task-relevant computations (Mastrogiuseppe and Ostojic 2018). In this vein, we sought to understand how the dynamics of RNNs could support the computations necessary for simulation-based M-Pong performance.

Figure 5A schematically illustrates the RNN hidden state space, with trajectories for two hypothetical trial conditions (dark blue, dark red) during the occluded epoch. We note that in order to accurately perform “simulation”, a projection of each trajectory must approximately correspond to the latent time-varying position of the ball [x,y] (light blue, light red, Figure 5A). During the visual epoch of the task, the position variables may be computed through nonlinear transformation of direct sensory input to the network. In contrast, during the occluded epoch when all sensory inputs have extinguished, RNNs behave as autonomous dynamical systems, and must therefore update the position based on a nonlinear function of network hidden states at the previous time steps. Since updating the position requires information about velocity, we reasoned that RNN states during the occluded epoch should additionally represent the ball velocity *[dx,dy]* (green subspace, Figure 5A).

**Figure 5:**
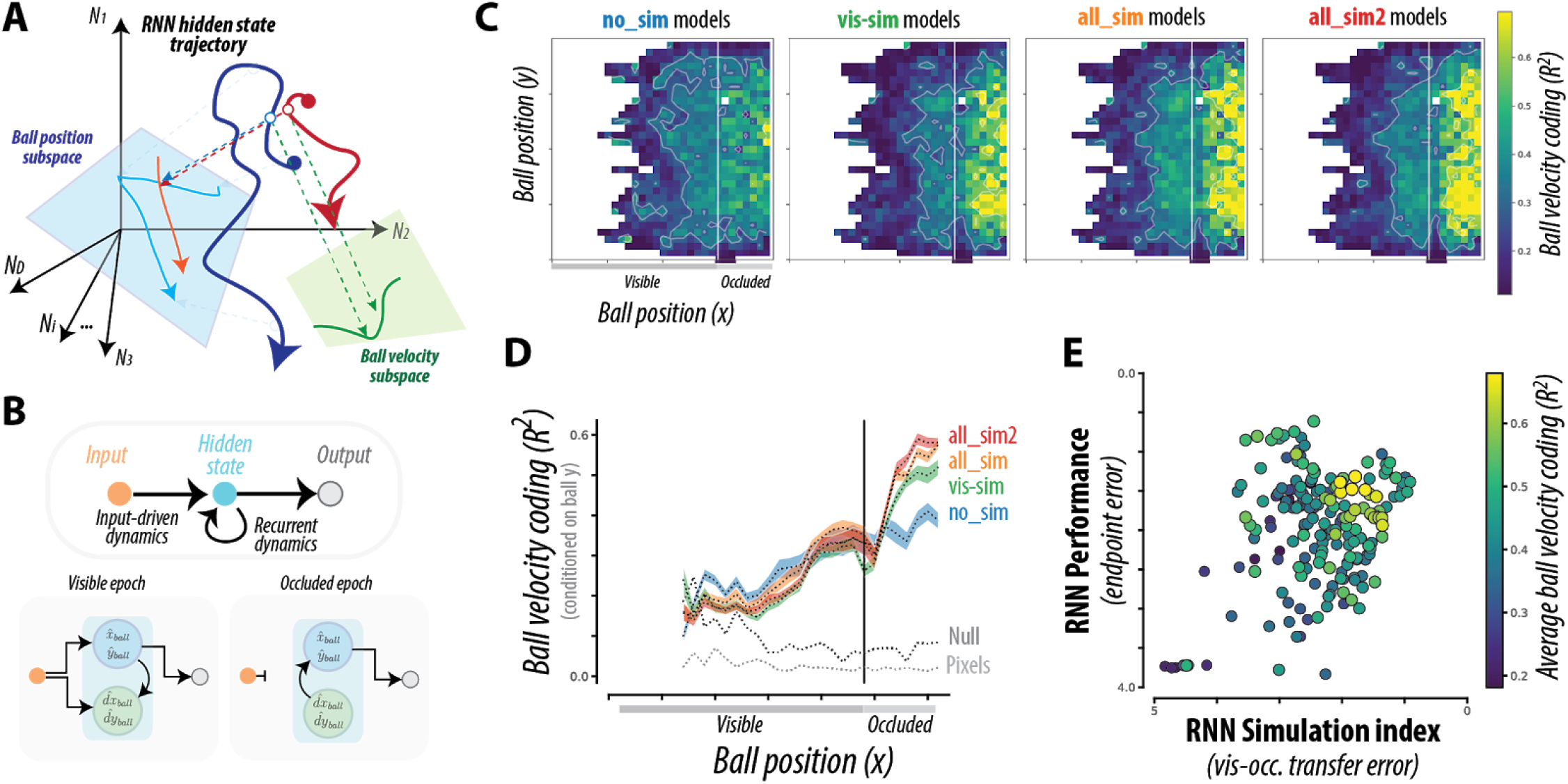
**(A) Conceptual schematic**. Schematic of RNN hidden state trajectory for two hypothetical conditions (dark blue, dark red) during the occluded epoch. To accurately perform “simulation”, a projection (illustrated in cyan) of each RNN hidden state trajectory must approximately correspond to the latent time-varying position of the ball [x,y] (here illustrated in light blue, light red). However, during the occluded epoch, RNNs are *autonomous* non-linear dynamical systems, wherein the hidden states evolve over time based on recurrent computations alone. Thus, we reasoned that RNN states additionally encode the ball velocity (green projection). **(B)** To produce different hidden state trajectories for different conditions, the RNN hidden state trajectories must be flexibly controlled by their initial conditions. Thus, we reasoned that RNNs must estimate both ball position and velocity from the sensory input during the visible epoch (left) and maintain ball velocity estimate throughout the occluded epoch to autonomously update the ball position (right). **(C)** Each panel shows a heat map of the M-Pong frame, with colors corresponding to the cross-validated proportion of variance of ball velocity explained (R2) by a linear read-out of the RNN hidden states, conditioned on ball position, and averaged over all RNNs of the same optimization type. **(D)** Ball velocity coding conditioned on the vertical ball position, averaged over all RNNs of the same optimization type. The dashed lines correspond to controls, verifying that velocity coding was not present in the sensory inputs (“pixels”), and not confounded with ball position (“null”). **(E)** Over all tested RNN models, the scatter shows the overall performance versus the simulation index, colored by the strength of the explicit representation of ball velocity during the occluded epoch. Ball velocity predictivity was greater for specific high-performing simulation-based RNN models, with strong correlations to both overall performance and simulation index. Note that the abscissa and ordinate are flipped such that left-to-right and bottom-to-top correspond with increasing performance (i.e., decreasing error) and increasing simulation ability (i.e. decreasing simulation error).

An autonomous RNN with no external input can behave differently for different trial conditions if and only if its initial condition at the beginning of the occluded epoch are appropriately adjusted (i.e., different initial conditions for different trial conditions). Accordingly, we reasoned that RNNs must use the visual epoch to establish an appropriate velocity-dependent initial condition for the subsequent occluded epoch. This hypothesis is schematically illustrated in Figure 5B, showing that RNNs estimate both ball position and velocity from the sensory input during the visible epoch (left) and maintain ball velocity estimate throughout the occluded epoch to autonomously update the ball position (right).

These considerations can be summarized in terms of a bipartite hypothesis that RNNs must use the visual information to extract velocity information during the visual epoch, and must have a representation of this velocity information during the occluded epoch. To test this hypothesis, we quantified the extent to which ball velocity is encoded in the hidden state representation of RNNs. Each panel in Figure 5C shows a heat map of the M-Pong frame, with colors corresponding to the cross-validated proportion of variance of ball velocity explained (*R^2^*) by a linear read-out of the RNN hidden states, averaged over all RNNs of the same optimization type (see Figure S5B-left for each individual RNN model, and Figure S5B-right for corresponding results using a single linear read-out). We observe the emergence of a representation of ball velocity during the visible segment, consistent with the first part of the hypothesis. Moreover, there was strong and persistent velocity coding at nearly all ball positions throughout the occluded segment, consistent with the second part of the hypothesis. This is further quantified in Figure 5D, which shows the ball velocity coding, conditioned on the vertical ball position. Moreover, we verified that such ball velocity coding was not present in the sensory inputs (“pixels” in Figure 5D), and not confounded with ball position (“null” in Figure 5D).

Interestingly, Figure 5C and 5D suggest an enhanced representation of ball velocity in RNN models that were directly optimized for simulation (e.g. *all_sim, all_sim2*). Indeed, across RNNs, explicit ball velocity coding during occlusion was correlated with both overall performance (*r = 0.57, p < 1e-17*) and simulation index (*r = 0.41, p < 1e-8*). This dependence was not observed for the corresponding ball velocity coding during the visible epoch (Figure S5C). This result is noteworthy as the networks that were optimized for simulation were only constrained to represent the position – not velocity.

Taken together, these results point to a critical role of the explicit representation of ball velocity in RNN hidden states, and suggest that establishing autonomous activity dynamics that approximate both ball position and velocity may be a key feature for neural networks to solve M-Pong using simulation-based strategies.

## Discussion

A hallmark of primate cognition is the ability to understand the causal structure of the physical world, and make general inferences that go far beyond the available sensory data. Cognitive neuroscience has advanced the influential hypothesis that these rich inference abilities depend critically on the mind’s ability to form internal models of the world and perform simulations over those models (Tenenbaum et al. 2011; Battaglia, Hamrick, and Tenenbaum 2013; Hegarty 2004; Hamrick 2019). Indeed, mental simulations are considered to be one of the key computational ingredients that enable humans to make inferences and predictions in the absence of sensory inputs.

There are however, critical open questions as to whether and when primates rely on mental simulations. First, the evidence in support of the mental simulation hypothesis is limited. The strongest and most recent evidence comes from studies that show a certain degree of consistency between human behavior and the behavior of high-level computer programs running simulations (Battaglia, Hamrick, and Tenenbaum 2013; Ullman et al. 2017). Whether such high-level computer programs are suitable abstractions for how neural systems compute is debatable (Ladenbauer et al. 2019). Second, while there has been significant progress in creating model-based neural network agents(Goodfellow et al. 2014; Higgins et al. 2016; Kulkarni et al. 2015), exerting flexible control over such neural models has proven challenging (Nalisnick et al. 2018). Third, somewhat paradoxically, model-free neural network agents that do not rely on mental simulations can outperform their model-based counterparts in rich environments such as Atari games (Hessel et al. 2017). Together, these considerations highlight the necessity of revisiting the mental simulation hypothesis using neural network models that afford flexible inferences.

To address this question, we designed a behavioral task that requires subjects – whether human, monkey, or artificial model – to infer the position of a ball moving across a two-dimensional frame with reflecting walls, in the face of occlusion. A plausible, but not necessary, strategy to solve this task is to mentally simulate the ball as it moves behind the occluder, and use this online estimation to predict the position of the ball at the time of interception. We note that this corresponds to an “online” simulation process, i.e. one that is synchronous with the dynamics of an unobserved external process, which may be computationally distinct from “offline” or asynchronous simulation processes thought to be relevant for imagination (Hassabis et al., 2007; Shepard & Metzler, 1971) or counterfactual reasoning (Gerstenberg & Tenenbaum, 2017).

In addition to capturing key computational demands of flexible physical inferences, the M-Pong task also exhibits a powerful feature: monkeys were able to rapidly learn the task and make highly flexible generalizations. Previous research on the neurobiology of physical inference and mental simulation has been largely limited to neuroimaging experiments in humans. Due to the inherent limitations of such non-invasive techniques, work in humans has only been able to delineate the neural basis of physical inferences at a macroscopic scale (Fischer et al. 2016; Zacks 2008). To gain a detailed understanding of the underlying neural circuits and mechanisms, it is important to establish a suitable animal model for mental simulation. Critically, a suitable animal model would not require over-training (with tens of thousands of repeats of the same stimulus-response contingencies) to capture this behavior, as this could generate alternative behavioral strategies (e.g. “automatized” stimulus-response policies or “memorization”) and corresponding spurious underlying neural strategies. To this end, monkeys have long served as an ideal model for human cognition. As such, the observations that monkeys can rapidly learn M-Pong – specifically, learn to make inferences in the face of occlusion within a single behavioral session, and generalize to novel conditions on the very first trial – is significant and can serve as a unique platform for future work to gain a detailed understanding of the neural and mechanisms of mental simulation in primates.

We compared primate behavior to RNNs that were either only trained to perform the task, or trained additionally to simulate the state of the ball. Networks that were not optimized for simulation were able to solve the task by finding an “automatized” nonlinear function that mapped sensory inputs to a suitable final paddle position. This finding corroborates recent advances in AI showing neural networks’ capacity to implement arbitrarily complex input-output mappings (Funahashi and Nakamura 1993; Collins, Sohl-Dickstein, and Sussillo 2016; Hammer 2000). Networks that were optimized for simulation were also able to attain primate-level performance, and by construction, carried an internal representation of ball position. Importantly however, the patterns of errors primates made while performing the task were highly structured and were only captured by RNNs that were additionally capable of simulating the ball position. This finding was not limited to specific parameterization of RNN models, but robust across hundreds of RNNs with different units, inputs, and architectures.

One desirable feature of RNN models is that they can be used to generate specific and testable hypotheses for the underlying computations in the brain (Sussillo and Barak 2013; Kanitscheider and Fiete 2017). To better understand the differences between the RNNs, we analyzed the structure of their internal state dynamics. We observed that the RNN state dynamics were generally simpler (i.e., slow and smooth) for RNNs that were optimized to simulate the ball position (Mastrogiuseppe and Ostojic 2018). This is unsurprising given that the ball position changed smoothly (Gao et al. 2017). Based on this observation and several recent reports of task-relevant slow dynamics in the primate brain(Wang et al. 2018; Sussillo et al. 2015; Mante et al. 2013), we wondered if the slow dynamics was the main factor for emulating primate-like behavior. To test this possibility, we analyzed the behavior of a new batch of RNNs that were optimized to perform the task and exhibit simple dynamics but not constrained to simulate the ball position. These networks were also able to perform the task but failed to capture primate behavior as well as RNNs optimized for simulation. Taken together, these results suggest that the primate brain solves the task by establishing slow dynamics that manifest an internal model of the ball position.

We further analyzed the internal dynamics of RNNs asking how the visual input early in the trial enables networks to simulate ball position later in the occluded portion of the trial. Since updating ball position depends on knowledge about velocity, we quantified the degree to which different RNNs carried information about velocity. The velocity information was stronger and more readily decodable (via linear decoders) in the subset of RNNs that were optimized for simulating ball position (Figure 4). This result highlights an additional signature of simulation in neural networks and serves as a prediction for future physiology experiments.

We note that RNN models presented here still lack details of biological neural networks (e.g. spiking activity, excitatory/inhibitory connections, and modular connectivity), and additional connectivity constraints need to be imposed to support the corresponding inferences (Michaels et al. 2019; Yang et al. 2019; Andalman et al. 2019). Similarly, RNNs presented here lack specific task details, e.g. RNNs did not receive sensory feedback about the position of the paddle throughout the trial but humans and monkeys did. As noted by previous work in sensorimotor control, this feedback could help reduce movement variability (Egger et al. 2019). However, RNNs in our work did not exhibit this type of output variability since computations implemented by the RNN units were not noisy. Moreover, with further analysis, we verified that RNNs did have access to an internal feedback signal related to the moment-by-moment position of the paddle through recurrent connections (Figure S5D). In other words, RNNs could indeed ‘see’ and make use of the paddle position through their internal dynamics. In spite of these implementation differences we were able to discover specific models that predict human behavior to near human-monkey consistency, suggesting that these models reflect specific task-relevant computations in the primate brain.

Our results build on the general framework of using machine learning approaches(LeCun, Bengio, and Hinton 2015) to create neural networks that successfully model behavior (Marblestone, Wayne, and Kording 2016; Richards et al. 2019; Storrs and Kriegeskorte 2019). However, unlike research in the sensory systems in which networks with superior performance are also superior in capturing behavior (Yamins et al. 2014; Kell et al. 2018), we found that network performance was not a good predictor of resemblance to primate behavior. Instead, the networks that most successfully capture primate behavior were those that were additionally constrained to perform simulations. This approach may be understood as a generalization of simultaneously optimizing on multiple tasks (Yang, Cole, and Rajan 2019; Yang et al. 2019), or of optimizing for specific tasks in the face of specific regularization (Sussillo et al. 2015; Lee and DiCarlo 2019), with the goal of building interpretable models of behavioral and neural phenomena (Saxe, Nelli, and Summerfield 2021). To this end, our work highlights a novel and general approach for testing hypotheses about specific inductive biases that govern human cognition by directly comparing models that do or do not implement those biases. We used this approach to test the role of simulations in making inferences, but the same logic can be applied to other hypothesized building blocks of cognition such as hierarchical information processing(Sarafyazd and Jazayeri 2019) and counterfactual reasoning(Hoch 1985).

## Methods

Two adult monkeys (Macaca mulatta; female), and twelve human subjects (18-65 years, gender not queried) participated in the experiments. The Committee of Animal Care and the Committee on the Use of Humans as Experimental Subjects at Massachusetts Institute of Technology approved the animal and human experiments, respectively. All procedures conformed to the guidelines of the National Institutes of Health.

### Behavioral task (M-Pong)

In *M-Pong*, the player controls the vertical position of a paddle along the right edge of the screen to intercept a ball as it moves rightward. On each trial, the ball starts at a random initial position (x0,y0) and a random initial velocity (dx_0_, dy_0_), and moves at a constant speed throughout the trial. The screen additionally contains a large rectangular occluder right before the interception point such that the ball’s trajectory is visible only during the first portion of the trial. Trial conditions were constrained by the following criteria: 1) the ball always moved rightward (dx>0), 2) the duration of the visible epoch was within a fixed range ([15,45] RNN timesteps or [624.9, 1874.7] ms), 3) the duration of the occluded epoch was within a fixed range ([15,45] RNN timesteps or [624.9, 1874.7] ms), 4) the number of times the ball bounced was within a fixed range ([0,1]). These constraints imposed some covariations between the ball parameters (e.g. trials where the ball started farther from the occluder tended to also have greater ball speed), as shown in Figure S1. We sampled up to 212480 unique conditions for RNN training, and 200 held-out conditions for testing RNNs, humans and monkeys. Stimuli and behavioral contingencies were controlled by an open-source software (MWorks; http://mworks-project.org/) running on an Apple Macintosh platform.

### Experimental procedures

#### Humans

Humans were seated in front of a computer in a dark room, under soft head restraint using a chin-rest. Stimuli were presented on a fronto-parallel 23-inch display (distance: approximately 67 cm; refresh rate: 60 Hz; resolution: 1920 by 1200) and behavioral responses were registered using a standard Apple keyboard. The subject’s eye position was tracked every 1 ms with an infrared camera (Eyelink 1000; SR Research Ltd, Ontario, Canada). Each trial initiated when the subject acquired and held gaze on a central fixation point (white circle, diameter: 0.5 degree in visual angle in size) within a window of 4 degree of visual angle for 200 ms. Following this fixation acquisition, subjects were allowed to make any eye movements and freely view the screen. Afterwards, the M-Pong condition was rendered onto the screen with the entire frame spanning 20 degrees of visual angle: the ball was rendered at its initial position (x0, y0), and the paddle was rendered in the central vertical position at the right edge of the frame. As shown in Figure 1C, the paddle was initially rendered as a small, transparent green square (0.5 deg x 0.5 deg), but turned into a full paddle (0.5 deg x 2.5deg) when the subject first initiated paddle movements (i.e. pressed a key). This feature enforced that subjects performed a movement (i.e. trigger the paddle) on all trials. For the remainder of the trial, subjects could freely view the monitor as the ball moved at its fixed velocity (dx0, dy0), and move the paddle up or down using a standard computer keyboard. The paddle position was updated on every screen refresh (i.e. every 16.6ms), and moved at a constant speed of 0.17deg/16ms = 0.01deg/ms. Trial ended when the ball reached the right-end of the screen. Upon the trial end, the occluder disappeared to give subjects feedback of their performance. If they successfully intercepted the ball, it would bounce off their paddle (see Figure 1C); if they had failed to intercept it, it would continue its path off the frame. Trials were separated by an inter-trial-interval of 750 ms.

In addition to the occluded condition, we also tested trials of the same M-Pong conditions under partial occlusion (opacity of occluder corresponding to 95%) where subjects could use visually-guided strategies to perform the task. Such visible trials were randomly interleaved on 25% of all trials. Data from visible trials were not included in the analyses presented in the main manuscript, but as expected, error on visible trials was significantly lower than on occluded trials (Figure S2A).

We collected behavioral data from 12 human subjects each performing 1 hour of M-Pong behavior. Each subject was tested on 50-100 unique task conditions (i.e., different initial ball position and velocity), in both visible and occluded conditions, all randomly interleaved. Trials from all 12 subjects were pooled together to characterize the human behavior over the complete dataset of 200 conditions. Behavioral error patterns were remarkably similar across subjects (Figure S2B). Altogether, we measured 8985 trials (6701 and 2284 under the occluded and visible conditions, respectively).

We additionally collected 2711 trials (2462 and 249 under the occluded and visible conditions, respectively) of behavioral data from two held-out human subjects performing the exact same conditions of the same task, with one small difference: the paddle size was not 0.5 x 2.5deg, but 0.5 x 1.75deg in size. For the purpose of the current work, this held-out data served as an independent validation of our human behavioral measurements, and were used to estimate a human-to-human consistency ceiling that could be directly compared to monkey-to-human consistency estimates.

#### Monkeys

Before the experiments, animals were implanted with three pins for head restraint using standard procedures (under general anesthesia and using sterile surgical techniques). During the experiments, animals were seated comfortably in a primate chair, and were head-restrained. For training purposes, we first acclimated animals to a 1 degree-of-freedom joystick placed right in front of the primate chair. Next, we started a curriculum for training animals to perform M-Pong. Animals were first trained to use the joystick to control the vertical position of a paddle, and then practiced M-Pong using 200 unique trial conditions. For all experiments, the stimuli were presented on a fronto-parallel 23-inch (58.4-cm) monitor at a refresh rate of 60 Hz. Similar to humans, animals’ eye position was tracked every 1 ms with an infrared camera (Eyelink 1000; SR Research Ltd, Ontario, Canada). The joystick voltage output (0-5V) was converted to one of three states (up: 3-5V; down: 0-2V; neutral: 2-3V), which was used to update the position of the paddle. The paddle position was updated in the exact same manner as was done in human experiments: the paddle position was updated on every screen refresh (i.e. every 16.6ms), and moved at a constant speed of 0.17deg/16ms = 0.01deg/ms.

We collected behavioral data over 32 behavioral sessions (monkey P: 12; monkey C: 20). Altogether, we measured 52837 trials (39472 and 13365 under the occluded and visible conditions, respectively) with even sampling from both monkeys (monkey P: 19788, 6716; monkey C: 19684, 6649; under occluded and visible conditions respectively). Behavioral error patterns were remarkably similar across the two monkeys (Figure S3B), and trials from both monkeys were pooled together to characterize monkey behavior.

### RNN optimization

We constructed different recurrent neural network (RNN) models optimized to perform the same task as humans and monkeys. We trained several hundred RNNs to map a series of visual inputs (pixel frames) to a movement output, where the target movement output corresponded to a prediction of the particular paddle position at a particular time point in order to intercept the ball. Different RNN models varied with respect to architectural parameters (different cell types, number of cells, regularization types, input representation types), and were differently optimized (one of four different target outputs, either with or without internal simulation). Critically, RNNs were not optimized to reproduce primate behavior, only to solve specific tasks. RNNs were trained using the TensorFlow 1.14 library using standard back-propagation and adaptive hyperparameter optimization techniques (Golub and Sussillo 2018); training each RNN took one to two days on a Tesla K20 GPU.

We trained two different RNN architectures: LSTM and GRU. We trained relatively small RNNs (10 or 20 cells) for relatively long durations (100-500 passes through the entire training set, or epochs). LSTM models had four different gates (input, input modulation, forget, and output gates), while GRU models had two different gates (reset and update gates). Each gate was parameterized by two weight matrices of size *N_input_ x N_cells_* and *N_cells_ x N_cells_* (where *N_input_=100*, and *N_cells_ = 10* or *20*). All RNNs parameters were initialized to zero prior to training.

Different classes of RNNs with different visual input representations were tested. Each unique task condition consisted of at most 90 timepoints or frames, and we rendered each frame of each trial as a 100×100 grayscale image. Note that we did not render the paddle, whose position is controlled by the output of the model. Given the relatively high dimensionality of this input data in the pixel space (90×10000 for each condition), we compressed it using two different encoding transformations. 1) We reduced the dimensionality of the data in pixel-space using principal components analysis, learning the PCA mapping iteratively using small batches of 32 trials at a time, from a subset of 512 total. 2) We additionally tested a higher-level visual representation based on 3-D Gabor wavelets, mimicking the output of neurons in area MT. This Gabor representation was computed with 3D convolutions of the down-sampled image stream with 16 3D Gabor wavelets with spatial and temporal frequencies matching prior work (Nishimoto and Gallant 2011). We reduced the dimensionality of the data in MT-like space using the same iterative PCA strategy.

Different classes of RNNs were trained with one of four different optimization types, which we code-named as no_sim, vis-sim, all_sim and all_sim2. All RNNs had the same number (7) of output channels, each one being an independent linear read-out from the RNN states. Depending on the optimization type, we varied how the overall loss was computed from these output channels (i.e. which output channels were included in estimating the loss). We defined the overall loss as the average of the time-averaged loss of each output channel, thus weighing each output channel equally. Figure 2C shows the optimization targets for each type. For all RNN types, one of the output channels (called “movement output”) corresponded to the paddle position, which was optimized to predict only two samples per trial: one consisting of the initial central paddle position, and the second corresponding to the particular paddle position at a particular time point in order to intercept the ball. As shown in Figure 2C, this was the only loss term for RNNs of the “no_sim” class. For the remaining RNNs, we additionally estimated a loss term from some of the other channels, as the mean squared error between the channel output and a target time-varying signal corresponding to the ball’s position (x,y) during specific trial epochs (“vis-sim”: visual epoch only; “all_sim”: entire trial; “all_sim2”: separate channels for visual and occluded epochs). This set of optimization choices explored different computational constraints regarding the specifics of “simulation.”

Each RNN architecture was trained with and without regularization on the output read-out weights, using either L1 or L2 norm loss. The weight of the regularization loss was measured at two different strengths (0.01, 0.1). Altogether, there were five possible regularization choices. While regularization generally helped with regards to both performance and human-consistency, we found no meaningful difference between L1 and L2 norm loss regularization.

To summarize, we constructed RNN models that varied with respect to several hyper-parameters: different cell types (rnn_type: LSTM or GRU), number of cells (n_hidden: 10 or 20), input representation types (input: pixel_pca or gabor_pca), and regularization types (reg: L1_0.01, L1_0.1, L2_0.01, or L2_0.1, or none); and were differently optimized (loss_weight_type: no_sim, vis_sim, all_sim, or all_sim2). Figure S3A,B shows the effect of each hyperparameter choice on performance metrics (task performance, simulation index) and primate consistency (with respect to both human and monkey behavior).

To investigate whether our results were robust to the extent of RNN training, we additionally tested the effect of the training dataset used for RNN optimization. To do so, we first selected RNN model architecture with the highest human-consistency score, and evaluated key RNN metrics (e.g. performance, simulation index, consistency to humans and to monkeys) while varying both the number of training epochs and the training data (number of training samples and distribution of training data). To test the latter, we first created a larger dataset of M-Pong trials with more variation in ball speed. We found that these metrics were largely insensitive to such variations in RNN optimization (see Figure S4A). Together, this suggests that the extent of RNN training was sufficient to converge upon “stable” network solutions, and that our key results and inferences are largely robust of the details of this optimization procedure.

Finally, we investigated whether the gains in human-consistency obtained via optimization for simulation could be explained by inducing slow and smooth dynamics in the RNN hidden states. To do so, we optimized a set of RNN models on task performance (as in “no_sim” models) but with additional regularization to promote slow and smooth dynamics, as in (Sussillo et al. 2015). Specifically, we added three regularization terms corresponding to the L2-norm of the hidden state activity, as the L2-norm of the derivative of the hidden state activity, and the ratio between these two. These three terms were weighted by three corresponding hyper-parameters, which we swept over a broad range in order to ensure that these regularizations had a significant effect on the learned RNN representations. Figure S5A shows the distribution of human-consistency as well as various representational metrics (dimensionality, speed, curvature, and norm) for all trained RNN models, grouped by their optimization type.

### RNN testing

With these trained RNNs in hand, we estimated a number of properties to characterize each model. Each trained RNN was tested on the same held-out set of 200 unique conditions that were tested in humans and monkeys. We measured the overall performance as the mean absolute error between the final model output (corresponding to a paddle position) and the ground truth final ball position. Given that not all networks were optimized to produce a read-out of the instantaneous ball position, we estimated their ability to “simulate” by training a linear read-out on the RNN states to predict the instantaneous ball position. This read-out was trained on both the visual and occluded epochs and tested on the occluded epoch, and we used a two-fold cross-validation scheme over conditions and time-points. We then quantified the simulation index as the mean absolute error between the predicted and true time-varying ball position.

### RNN characterization

We characterized each RNN’s internal state representation via a number of representational attributes. The state representation consists of a matrix ***X*** of size *N_trials_× N_timesteps_ × N_units_*. We first computed the first and second discrete temporal derivatives ***X’*** and ***X”***. We estimated normalized trajectory speed via the average absolute first derivative, normalized by the average absolute position. Similarly, we estimated a metric of trajectory curvature via the average absolute second derivative, normalized by the average absolute position. To mathematically define these metrics, we first define, for a matrix ***A*** of size *N_trials_× N_timesteps_ × N_units_*, the norm over units ‖*A*‖*_k_* and the average over timesteps μ(*A*)*j* as:

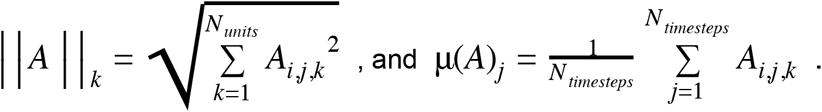

Using this notation, the normalized trajectory speed and curvature metrics correspond to:

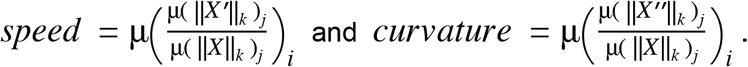

We then reshaped the state representation matrix of each RNN, concatenating the trial and timestep dimensions, into **X_mat_** of size *N_trials × timesteps_ × N_units_*. We estimated the dimensionality of **X_mat_** over all conditions and over the entire trial via a participation ratio metric, a weighted sum of the eigen-spectrum obtained from PCA.

For each RNN model, we additionally estimated a measure of “feedback control” to characterize the alignment between the read-out weights and the recurrent weights. While RNNs do not receive explicit instantaneous visual feedback, this metric aims to capture the extent to which the output of the network is fed back into its activity. For each trained RNN model and for each RNN gate type, we extracted the matrices of input weights (*N_input_D_ × N_units_*), recurrent weights (*N_units_ × N_units_*), and read-out weights (*Nunits × 1*).

We then computed a metric of feedback control as the normalized projection of the read-out weights onto each of the recurrent weights (i.e. the dot product of the corresponding unit vectors in weight space), and averaging across weights and gate types. Note that our LSTM models have four different gates (*input*, *input modulation*, *forget*, and *output* gates), while GRU models have two different gates (*reset* and *update* gates). We observed that the median amount of feedback control was significantly greater than that expected from random read-out weights, across RNNs (Figure S5D).

### RNN analysis

We note that the different RNN optimization types do not correspond to mutually exclusive hypotheses, but instead map on to overlapping parts of the hypothesis space, as shown in Figure 3A. For instance, the set of RNNs that were optimized for both task performance and simulation (e.g. “all_sim” and “all_sim2”) form a subset of the set of RNNs optimized on task performance alone (“no_sim”). Similarly, the “all_sim” RNNs form a subset of the set of RNNs optimized on task performance and simple dynamics (“simple_dynamics”). As a result, RNNs constructed from different optimization types were not explicitly required to differ with respect to their attributes. For example, RNNs optimized for simulation ability only during the visible epoch (“vis-sim”) could still exhibit strong simulation ability during the occluded epoch, despite not being directly optimized for this characteristic.

Given this potential for overlap between optimization types, we did not focus our analyses on group comparisons of consistency scores between the different optimization types. Instead, we sought to infer whether consistency scores depended on specific RNN attributes, over all RNN models, irrespective of the underlying optimization type. We quantified the strength of this dependence via the proportion of variance (*R^2^*) of consistency scores that could be explained by each RNN attribute. Moreover, to account for possible co-variations between RNN attributes and infer the proportion of variance that could be uniquely explained by each RNN attribute, we estimated partial *R^2^* (see Comparison metrics).

### Behavioral metrics

We first quantified a grand average estimate of performance using the mean absolute error (MAE), computed as the absolute difference between the final ball position and the final paddle position, averaged across all trials and all conditions.

To go beyond the summary statistic of global performance, we characterized primate and model behavior on this task using a pattern of errors across conditions. This process consists of mapping the final paddle position from a set of trials X (*N_trails_ x 1*) and the corresponding ground truth paddle positions Z (*N_trails_ x1*) to a pattern of errors Ymu (*N_cond_ x 1*, where *N_cond_=200*). However, we cannot simply measure the error of each trial *i* as a difference between the final paddle position Xi and the corresponding ground truth position Zi, as this will introduce spurious correlations between otherwise unrelated datasets X_1_ and X_2_ (e.g. correlations between X_1_-Z and X_2_-Z). To mitigate this, we first computed error patterns as residuals from a linear least squares regression Y = X – X_pred_, where X_pred_ = m*Z+b is the linear least squares fit of X. We then averaged trials of the same condition to obtain the pattern of residuals across conditions Y_mu_. Note that, with this definition of error, Pearson correlations between metrics estimated from two datasets is equivalent to computing a partial correlation between the pattern of endpoint paddle positions across conditions, accounting for the co-varying pattern of ground truth positions.

### Behavioral consistency

To quantify the similarity between humans and a model with respect to a given behavioral metric, we used a measure called the “*human-consistency*” 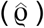 as previously defined (Johnson, Hsiao, and Yoshioka 2002). *Human-consistency* is computed as a noise-adjusted correlation of behavioral patterns (DiCarlo and Johnson 1999; Spearman 1961). For each system (model or human), we randomly split all behavioral trials into two equal halves and estimated the behavioral pattern from each half, resulting in two independent estimates of the system’s behavioral pattern. We took the Pearson correlation between these two estimates as a measure of the reliability of that behavioral pattern given the amount of data collected, i.e. the split-half internal reliability. To estimate the *human-consistency*, we computed the Pearson correlation over all the independent estimates of the behavioral pattern from the model (**m**) and the human (**h**), and we then divide that raw Pearson correlation by the geometric mean of the split-half internal reliability of the same behavioral pattern measured for each system:

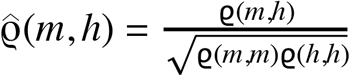

Since all correlations in the numerator and denominator were computed using the same amount of trial data (exactly half of the trial data), we did not need to make use of any prediction formulas (e.g. extrapolation to larger number of trials using Spearman-Brown prediction formula). This procedure was repeated 10 times with different random split-halves of trials. Our rationale for using a reliability-adjusted correlation measure for *human-consistency* was to account for variance in the behavioral signatures that arises from “noise,” i.e., variability that is not replicable by the experimental condition (image and task) and thus that no model can be expected to predict. In sum, if the model (**m**) is a replica of the archetypal human (**h**), then its expected human-consistency is 1.0, regardless of the finite amount of data that is collected.

### Comparison metrics

In addition to reporting individual model scores with respect to this behavioral benchmark (Schrimpf et al. 2018), we investigated what specific attributes of models best predicts their human-consistency. We estimated the relative importance of specific attributes using a Pearson correlation. To account for covariations due to other attributes, we also report a partial Pearson correlation, the estimated correlation after regressing out co-varying attributes with a linear least squares regression.

For several analyses, we measured the proportion of variance explained of a high-dimensional signal X by another high-dimensional signal Y using an *R^2^* metric. To estimate *R^2^* in an unbiased manner, we performed the following analysis. We first orthogonalized the matrix X into X’ using a PCA preprocessing step. We used 5-fold cross-validation to run a linear regression to predict each dimension of X’ from Y. We estimated the proportion of variance as the square of the Pearson correlation R^2^ from this prediction. Across columns of X’, this resulted in a vector of *R^2^* for each dimension; we measured the total variance explained as a weighted sum of this vector, with weights corresponding to the eigenvalues of the covariance matrix of X (i.e. the proportion of variance of X explained by each dimension of X’).

## Acknowledgments

R.R. is supported by the Helen Hay Whitney Foundation. M.J. is supported by NIH (NIMH-MH122025), the Simons Foundation, the McKnight Foundation, and the McGovern Institute.

