## Supplemental Materials for "The role of mental simulation in primate physical inference abilities"

**Supplemental Information**

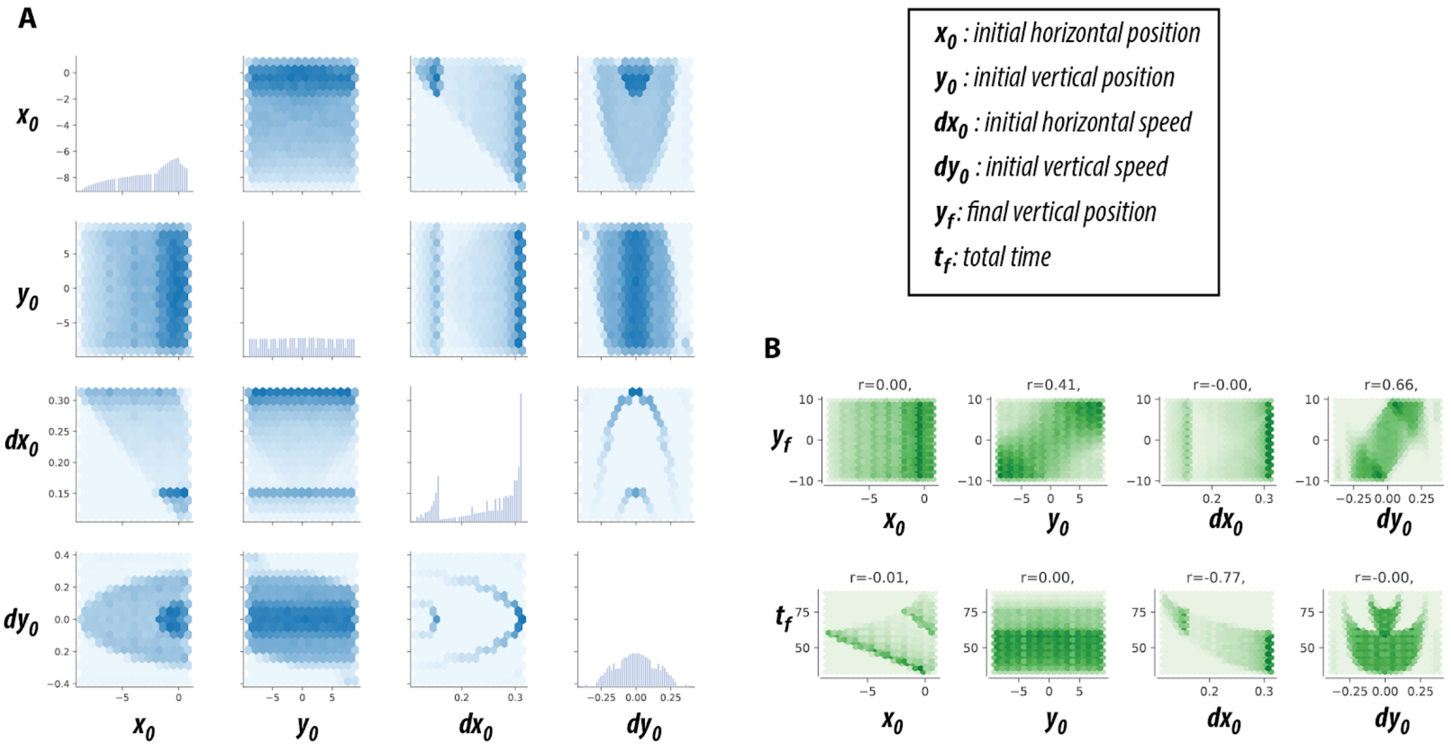


**Figure S1. Task parameters.** **(A)** Distribution of meta-parameters corresponding to initial position ($x_0$,$y_0$) and initial velocity ($dx_0$, $dy_0$) for the task RNN training set. The diagonal panels correspond to histograms of each meta-parameter. The off-diagonal panels correspond to 2D histograms showing the joint distribution for each pair of meta-parameters. **(B)** We sampled the initial ball position and velocity sufficiently broadly enough to ensure that the task was reasonably challenging. Each panel shows the joint distribution of each meta-parameter with the target output (final ball position, $y_f$, see top panels) and the time to interception ($t_f$, see bottom panels).

**Figure S2:**

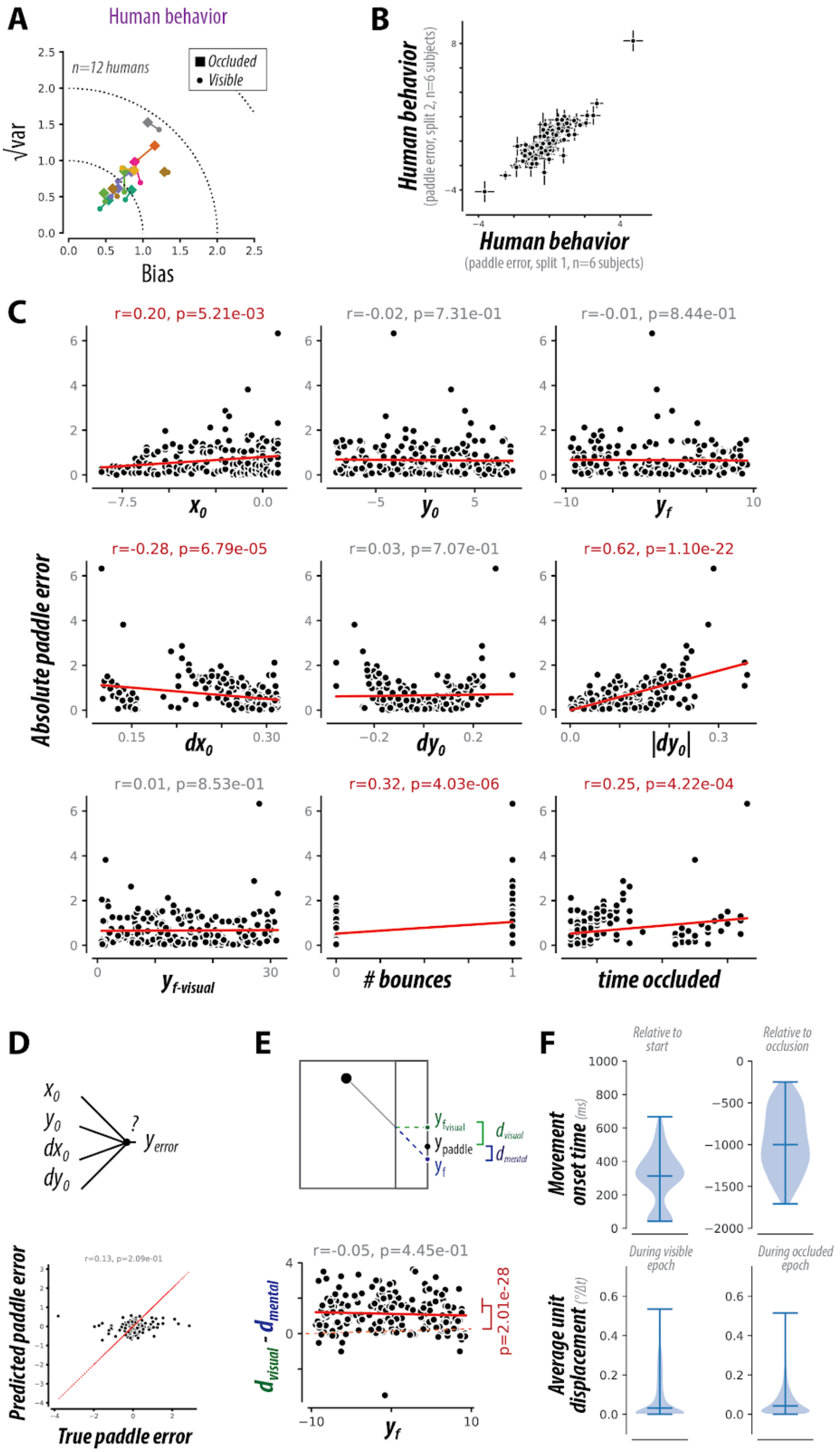
A
Human behavior
n=12 humans
Occluded
Visible
√var
Bias
B
Human behavior
(paddle error, split 2, n=6 subjects)
Human behavior
(paddle error, split 1, n=6 subjects)
C
r=0.20, p=5.21e-03
r=-0.02, p=7.31e-01
r=-0.01, p=8.44e-01
$x_0$
$y_0$
$y_f$
r=-0.28, p=6.79e-05
r=0.03, p=7.07e-01
r=0.62, p=1.10e-22
Absolute paddle error
$dx_0$
$dy_0$
$|dy_0|$
r=0.01, p=8.53e-01
r=0.32, p=4.03e-06
r=0.25, p=4.22e-04
$y_{f\text{-visual}}$
### bounces
time occluded
D
$x_0$
$y_0$
$dx_0$
$dy_0$
?
$y_{error}$
r=0.13, p=2.09e-01
Predicted paddle error
True paddle error
E
$y_{f\text{visual}}$
$y_{\text{paddle}}$
$y_f$
$d_{visual}$
$d_{mental}$
r=-0.05, p=4.45e-01
$d_{visual} - d_{mental}$
p=2.01e-28
$y_f$
F
Relative to start
Relative to occlusion
Movement onset time (ms)
During visible epoch
During occluded epoch
Average unit displacement (°/Δt)

**(A)** Human behavior for each individual subject, separately for visible and occluded trials. Recall that visible trials, which had a partially opaque occluder, were randomly interleaved with occluded trials. Performance is shown as a scatter of the bias (absolute error, after averaging across trials of the same condition) and variance (variability across trials of the same condition). The dotted circles correspond to lines of equal root-mean-squared error (RMSE). As expected, error is lower on visible trials.

**(B)** Comparison of behavioral error patterns between two spit-halves of human subjects reveals virtually identical error patterns, demonstrating that we measured a highly reliable human behavioral pattern. Trials from all 12 subjects were pooled to characterize the “archetypal” human behavior.

**(C)** Dependence of error patterns on task variables. Each panel shows absolute average endpoint error against task parameters varying per condition. The strength of each dependence, measured via a Pearson correlation, is shown on the corresponding panel titles; significant dependences are highlighted in red. Importantly, error patterns depended largely on dynamic variables relating to the ball speed ($dx_0$, $dy_0$, $|dy_0|$, time_occluded) but not initial, intermediate, or final ball position ($y_0$, $y_{f\_visual}$, $y_f$). Given the correlation between initial x position and ball speed built into this dataset (see Figure S1), errors were additionally correlated to the initial x position ($x_0$). The occurrence of bounces also caused significantly greater errors, on average.

**(D)** Error patterns could not be explained by a simple function of initial position and velocity. We used cross-validated linear regression to predict the error pattern from the initial ball position and ball velocity; the resulting prediction was not better than expected by chance.

**(E)** Error patterns could not be explained by a simple visual tracking strategy, where the final paddle position is estimated based on the last visible position of the ball ($y_{f\_visible}$). To demonstrate this, we compared the distance between the average final paddle position to each of the candidate final positions, as predicted by visual tracking ($d_{visual}$) and mental tracking ($d_{mental}$) strategies. We observe that the corresponding bias ($d_{visual}$ - $d_{mental}$) is significantly greater than zero (see red annotation) and moreover is not dependent on the final ball position (as might be expected if this bias is somehow driven by boundary conditions, see gray title annotation), demonstrating that behavioral error patterns are inconsistent with a simpler visual tracking strategy.

**(F)** Dynamics of movements. The top panels show the distribution of movement onset times, relative to the beginning of the trial (left) and to the beginning of the occluded epoch (right). Subjects initiated movements ~1000ms before the beginning of the occluded epoch. The bottom panels show the distribution of unit displacement during the visible and occluded epochs. Unit displacement was estimated from the instantaneous paddle position by first averaging across trials of the same conditions, and then measuring the mean absolute change in position. Subjects moved the paddle in both visible and occluded epochs, but displacements were significantly larger in the occluded epoch.

**Figure S3:**

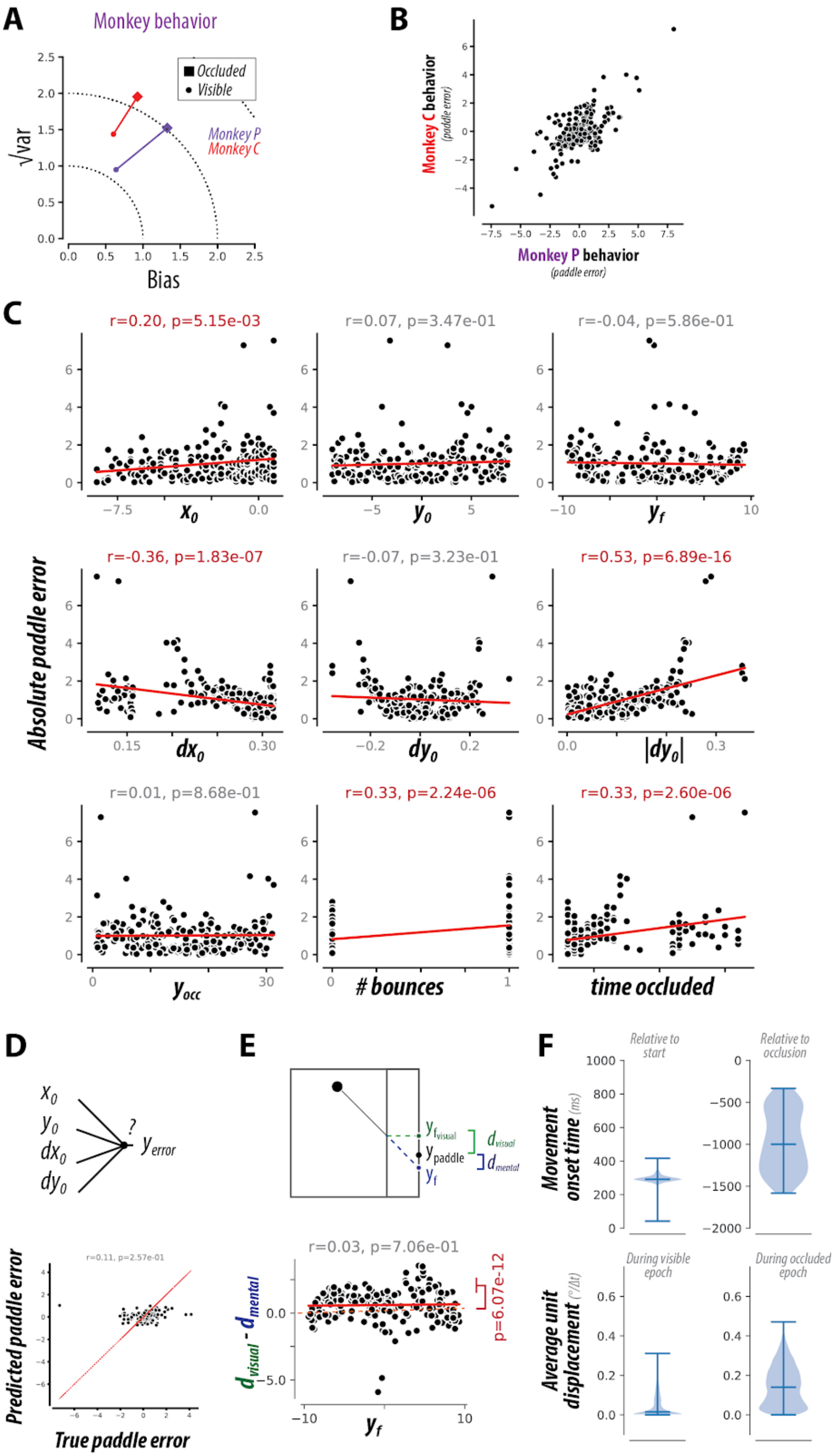


**(A-F)** Formatting identical to Figure S2, but for monkey behavior.

**Figure S4:**

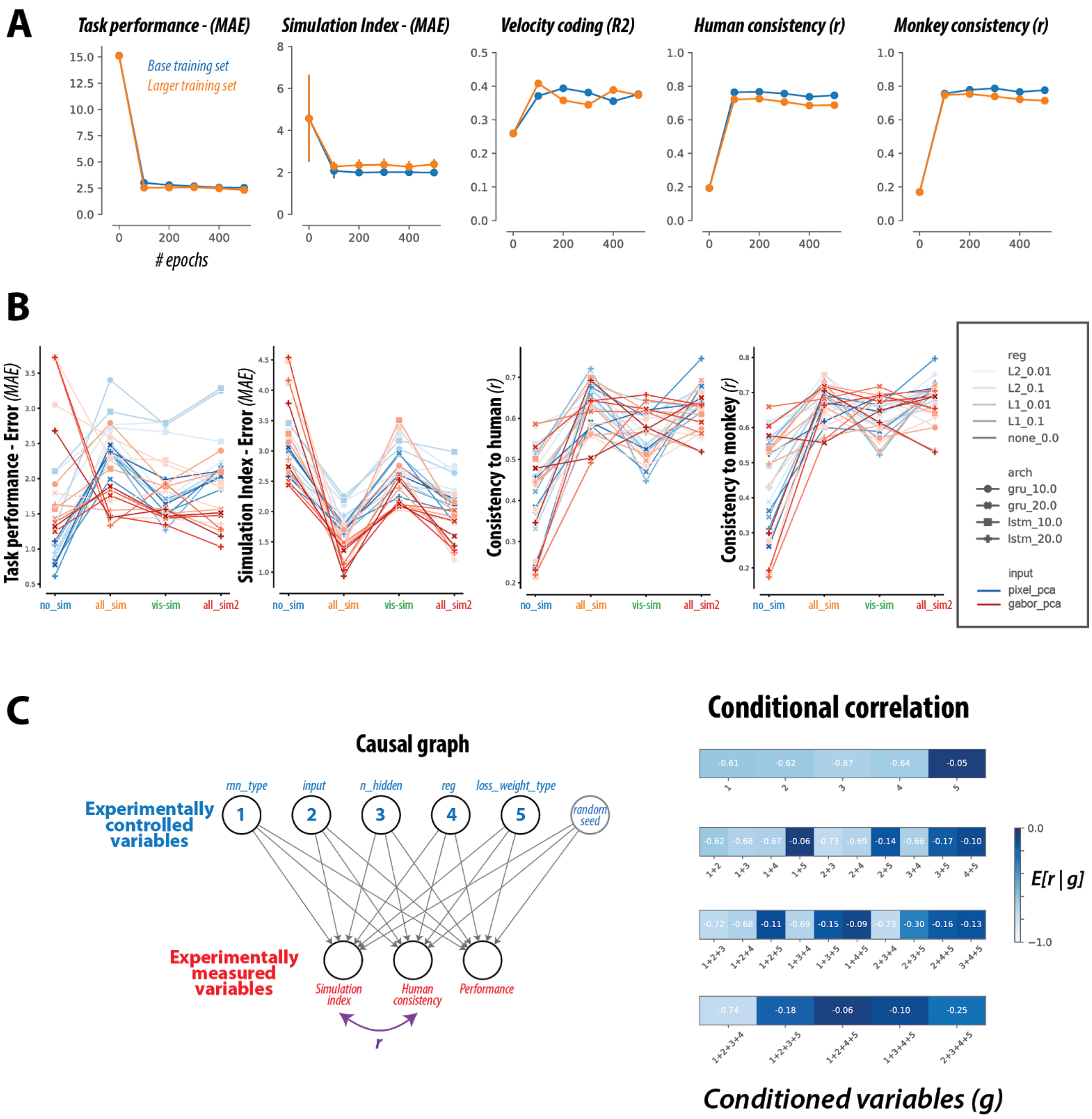


**(A)** Effect of RNN optimization. For the RNN model architecture with the highest human-consistency score, we evaluated key RNN metrics (e.g. performance, simulation index, consistency to humans and to monkeys) while varying both the number of training epochs and the training data (number of training samples and distribution of training data). We found that these metrics were largely insensitive to such variations in RNN optimization, suggesting that the extent of RNN training was sufficient to converge upon "stable" network solutions, and that our key results and inferences are largely robust of the details of this optimization procedure. **(B)** Different RNN models varied with respect to several hyper-parameters: different cell types (***rnn_type***: *LSTM or GRU*), number of cells (***n_hidden***: *10 or 20*), input representation types (***input***: *pixel_pca or gabor_pca*), and regularization types (***reg***: *L1_0.01, L1_0.1, L2_0.01, or*

*L2_0.1*); and were differently optimized (***loss_weight_type****: no_sim, vis_sim, all_sim, or all_sim2*). Each of the four panels shows the effect of each hyperparameter choice on performance metrics (task performance, simulation index) and primate consistency (with respect to both human and monkey behavior). **(B)** The left panel shows the causal graph of our RNN experiments. We experimentally controlled five different RNN hyper-parameters (top row, blue), and from the resulting RNN model instances, we measured several attributes (bottom row, red), including the simulation index and the human consistency. To uncover which RNN hyper-parameters *cause* the strong negative correlation (r) between simulation index and human consistency, we measured the conditional correlation E[r|g], conditioning on each of the 30 possible combinations (g) of the five hyper-parameter types. The right panel shows the resulting conditional correlations for each of the 30 combinations. Darker values correspond to smaller magnitude correlations. The observed correlation between the simulation index and human consistency is largely driven by the hyper-parameter defining the optimization target (***loss_weight_type***).

**Figure S5:**

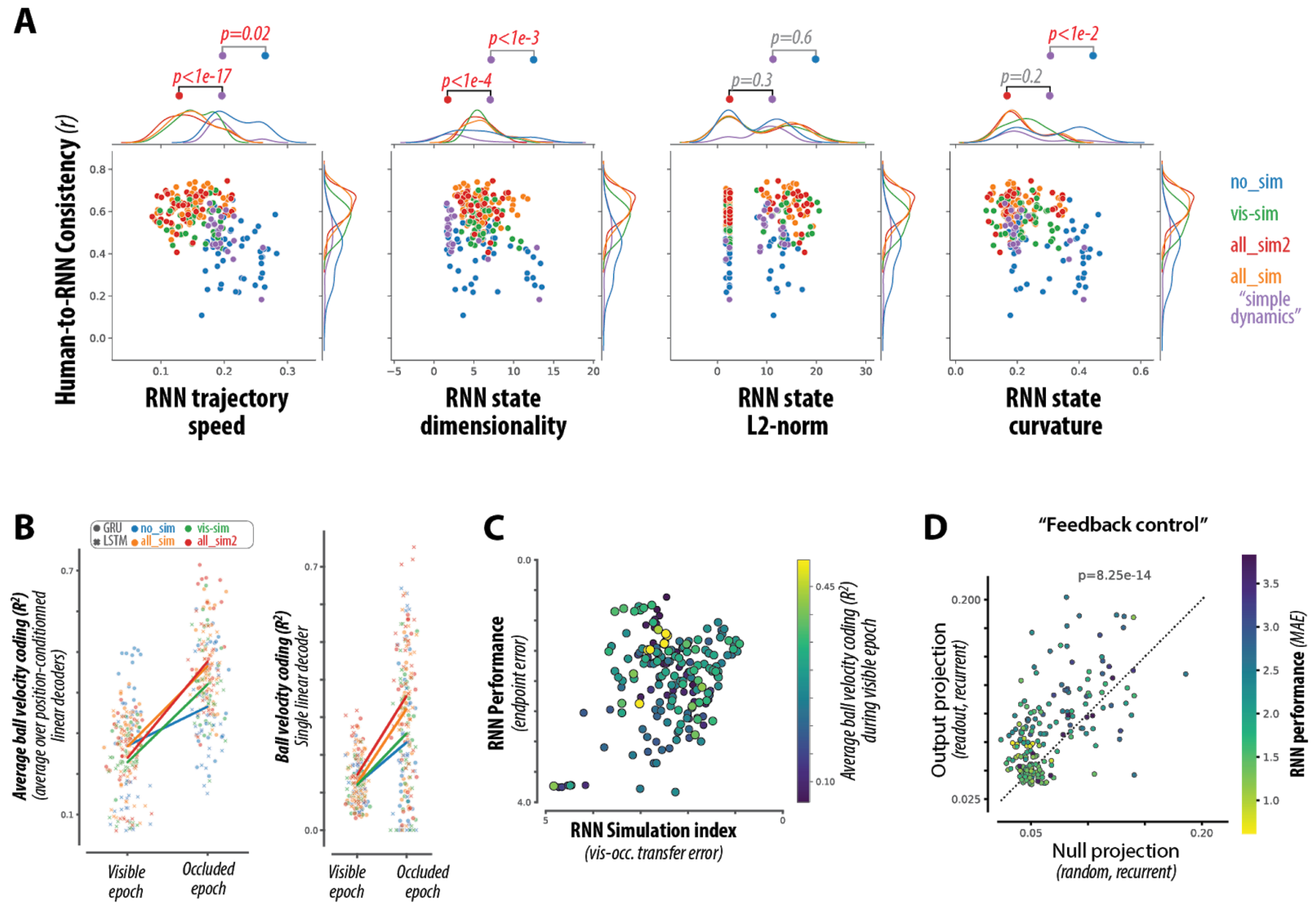


**(A)** As a control, we optimized a new set of RNN models on task performance with additional regularization to promote simple dynamics, by adding regularization terms related to the L2-norms of the hidden state activity and the derivative of the hidden state activity (see Methods). To verify that this regularization had the intended effect on RNN representations, the four panels show the distribution of human-consistency against four representational metrics (dimensionality, speed, curvature, and norm) for all trained RNN models, grouped by their optimization type. Statistical comparisons between relevant distributions are shown above each scatter (unpaired t-test). The top comparison corresponds to 'no_sim' vs 'simple_dynamics' (blue vs purple), and the bottom comparison corresponds to 'all_sim2' vs 'simple_dynamics' (red vs purple). We observe that for all metrics that differed across RNN types, the 'simple_dynamics' RNNs (purple) did indeed diverge from the baseline 'no_sim' (blue) models as intended. **(B)** Comparison of average velocity coding during visible and occluded epochs, for all RNN models. The left panel shows velocity coding estimated via position-conditioned linear decoders, whereas the right panel shows the corresponding estimates with a single position-independent linear decoder. **(C)** Average velocity coding during visible epoch is not correlated to either performance or simulation index, in contrast to the corresponding metric during the occluded epoch (see Figure 5E). **(D)** We estimated a measure of "feedback control" to characterize the alignment between the read-out weights and the recurrent weights. While RNNs do not receive explicit instantaneous visual feedback, this metric aims to capture the extent to which the output of the network is fed back into its activity. The scatter plot shows the comparison of this metric against the null, for each RNN model; marker color corresponds to model performance. We observed that the median amount of feedback control was significantly greater than expected by chance.

Hoch, Stephen J. 1985. "Counterfactual Reasoning and Accuracy in Predicting Personal Events." *Journal of*

*Experimental Psychology. Learning, Memory, and Cognition* 11 (4): 719–31.
Johnson, Kenneth O., Steven S. Hsiao, and Takashi Yoshioka. 2002. “Neural Coding and the Basic Law of Psychophysics.” *The Neuroscientist: A Review Journal Bringing Neurobiology, Neurology and Psychiatry* 8 (2): 111–21.
Kanitscheider, Ingmar, and Ila Fiete. 2017. “Training Recurrent Networks to Generate Hypotheses about How the Brain Solves Hard Navigation Problems.” In *Advances in Neural Information Processing Systems*, edited by I. Guyon, U. V. Luxburg, S. Bengio, H. Wallach, R. Fergus, S. Vishwanathan, and R. Garnett, 30:4529–38. Curran Associates, Inc.
Kell, A. J. E., Daniel L. K. Yamins, Erica N. Shook, Sam V. Norman-Haignere, and Josh H. McDermott. 2018. “A Task-Optimized Neural Network Replicates Human Auditory Behavior, Predicts Brain Responses, and Reveals a Cortical Processing Hierarchy.” *Neuron* 98 (3): 630–44.e16.
Kulkarni, Tejas D., William F. Whitney, Pushmeet Kohli, and Josh Tenenbaum. 2015. “Deep Convolutional Inverse Graphics Network.” In *Advances in Neural Information Processing Systems*, edited by C. Cortes, N. Lawrence, D. Lee, M. Sugiyama, and R. Garnett, 28:2539–47. Curran Associates, Inc.
Ladenbauer, Josef, Sam McKenzie, Daniel Fine English, Olivier Hagens, and Srdjan Ostojic. 2019. “Inferring and Validating Mechanistic Models of Neural Microcircuits Based on Spike-Train Data.” *Nature Communications* 10 (1): 4933.
LeCun, Yann, Yoshua Bengio, and Geoffrey Hinton. 2015. “Deep Learning.” *Nature* 521 (7553): 436–44.
Lee, Hyodong, and James J. DiCarlo. 2019. “Topographic Deep Artificial Neural Networks (TDANNs) Predict Face Selectivity Topography in Primate Inferior Temporal (IT) Cortex.” *arXiv [q-bio.NC]*. arXiv. http://arxiv.org/abs/1909.09847.
Lerer, Adam, Sam Gross, and Rob Fergus. 2016. “Learning Physical Intuition of Block Towers by Example.” *arXiv [cs.AI]*. arXiv. http://arxiv.org/abs/1603.01312.
Maheswaranathan, N., A. Williams, and M. Golub. 2019. “Universality and Individuality in Neural Dynamics across Large Populations of Recurrent Networks.” *Advances in Neural Information Processing Systems*. http://papers.nips.cc/paper/9694-universality-and-individuality-in-neural-dynamics-across-large-populations-of-recurrent-networks.
Mante, Valerio, David Sussillo, Krishna V. Shenoy, and William T. Newsome. 2013. “Context-Dependent Computation by Recurrent Dynamics in Prefrontal Cortex.” *Nature* 503 (7474): 78–84.
Marblestone, Adam H., Greg Wayne, and Konrad P. Kording. 2016. “Toward an Integration of Deep Learning and Neuroscience.” *Frontiers in Computational Neuroscience* 10 (September): 94.
Mastrogiuseppe, Francesca, and Srdjan Ostojic. 2018. “Linking Connectivity, Dynamics, and Computations in Low-Rank Recurrent Neural Networks.” *Neuron* 99 (3): 609–23.e29.
Michaels, Jonathan A., Benjamin Dann, and Hansjörg Scherberger. 2016. “Neural Population Dynamics during Reaching Are Better Explained by a Dynamical System than Representational Tuning.” *PLoS Computational Biology* 12 (11): e1005175.
Michaels, Jonathan A., Stefan Schaffelhofer, Andres Agudelo-Toro, and Hansjörg Scherberger. 2019. “A Neural Network Model of Flexible Grasp Movement Generation.” https://doi.org/10.1101/742189.
Nalisnick, Eric, Akihiro Matsukawa, Yee Whye Teh, Dilan Gorur, and Balaji Lakshminarayanan. 2018. “Do Deep Generative Models Know What They Don’t Know?” *arXiv [stat.ML]*. arXiv. http://arxiv.org/abs/1810.09136.
Nishimoto, Shinji, and Jack L. Gallant. 2011. “A Three-Dimensional Spatiotemporal Receptive Field Model Explains Responses of Area MT Neurons to Naturalistic Movies.” *The Journal of Neuroscience: The Official Journal of the Society for Neuroscience* 31 (41): 14551–64.
Remington, Evan D., Devika Narain, Eghbal A. Hosseini, and Mehrdad Jazayeri. 2018. “Flexible Sensorimotor Computations through Rapid Reconfiguration of Cortical Dynamics.” *Neuron* 98 (5): 1005–19.e5.
Richards, Blake A., Timothy P. Lillicrap, Philippe Beaudoin, Yoshua Bengio, Rafal Bogacz, Amelia Christensen, Claudia Clopath, et al. 2019. “A Deep Learning Framework for Neuroscience.” *Nature Neuroscience* 22 (11): 1761–70.
Russo, Abigail A., Sean R. Bittner, Sean M. Perkins, Jeffrey S. Seely, Brian M. London, Antonio H. Lara, Andrew Miri, et al. 2018. “Motor Cortex Embeds Muscle-like Commands in an Untangled Population Response.” *Neuron* 97 (4): 953–66.e8.
Sarafyazd, Morteza, and Mehrdad Jazayeri. 2019. “Hierarchical Reasoning by Neural Circuits in the Frontal

Cortex.” *Science* 364 (6441). https://doi.org/10.1126/science.aav8911.
Schrimpf, Martin, Jonas Kubilius, Ha Hong, Najib J. Majaj, Rishi Rajalingham, Elias B. Issa, Kohitij Kar, et al. 2018. “Brain-Score: Which Artificial Neural Network for Object Recognition Is Most Brain-Like?” *bioRxiv*. https://doi.org/10.1101/407007.
Shepard, R. N., and J. Metzler. 1971. “Mental Rotation of Three-Dimensional Objects.” *Science* 171 (3972): 701–3.
Sohn, Hansem, Devika Narain, Nicolas Meirhaeghe, and Mehrdad Jazayeri. 2019. “Bayesian Computation through Cortical Latent Dynamics.” *Neuron* 103 (5): 934–47.e5.
Spearman, Charles. 1961. “The Proof and Measurement of Association between Two Things.” https://psycnet.apa.org/record/2006-10257-005.
Storrs, Katherine R., and Nikolaus Kriegeskorte. 2019. “Deep Learning for Cognitive Neuroscience.” *arXiv [q-bio.NC]*. arXiv. http://arxiv.org/abs/1903.01458.
Sussillo, David, and Omri Barak. 2013. “Opening the Black Box: Low-Dimensional Dynamics in High-Dimensional Recurrent Neural Networks.” *Neural Computation* 25 (3): 626–49.
Sussillo, David, Mark M. Churchland, Matthew T. Kaufman, and Krishna V. Shenoy. 2015. “A Neural Network That Finds a Naturalistic Solution for the Production of Muscle Activity.” *Nature Neuroscience* 18 (7): 1025–33.
Tenenbaum, Joshua B., Charles Kemp, Thomas L. Griffiths, and Noah D. Goodman. 2011. “How to Grow a Mind: Statistics, Structure, and Abstraction.” *Science* 331 (6022): 1279–85.
Ullman, Tomer D., Elizabeth Spelke, Peter Battaglia, and Joshua B. Tenenbaum. 2017. “Mind Games: Game Engines as an Architecture for Intuitive Physics.” *Trends in Cognitive Sciences* 21 (9): 649–65.
Wang, Jing, Devika Narain, Eghbal A. Hosseini, and Mehrdad Jazayeri. 2018. “Flexible Timing by Temporal Scaling of Cortical Responses.” *Nature Neuroscience* 21 (1): 102–10.
Yamins, Daniel L. K., Ha Hong, Charles F. Cadieu, Ethan A. Solomon, Darren Seibert, and James J. DiCarlo. 2014. “Performance-Optimized Hierarchical Models Predict Neural Responses in Higher Visual Cortex.” *Proceedings of the National Academy of Sciences of the United States of America* 111 (23): 8619–24.
Yang, Guangyu Robert, Michael W. Cole, and Kanaka Rajan. 2019. “How to Study the Neural Mechanisms of Multiple Tasks.” *Current Opinion in Behavioral Sciences* 29 (October): 134–43.
Yang, Guangyu Robert, Madhura R. Joglekar, H. Francis Song, William T. Newsome, and Xiao-Jing Wang. 2019. “Task Representations in Neural Networks Trained to Perform Many Cognitive Tasks.” *Nature Neuroscience* 22 (2): 297–306.
Zacks, Jeffrey M. 2008. “Neuroimaging Studies of Mental Rotation: A Meta-Analysis and Review.” *Journal of Cognitive Neuroscience* 20 (1): 1–19.
